# An environmentally responsive transcriptional state modulates cell identities during root development

**DOI:** 10.1101/2022.03.04.483008

**Authors:** Marina Oliva, Tim Stuart, Dave Tang, Jahnvi Pflueger, Daniel Poppe, Jafar S. Jabbari, Scott Gigante, Jonathan Michael Dragwidge, James Whelan, Mathew G. Lewsey, Ryan Lister

**Author notes:** **Correspondence:** Ryan Lister, Marina Oliva.

## Abstract

Roots are fundamental organs for plant development and response to their environment: they anchor the plant to its growth substrate, uptake nutrients and water vital to plant growth, and can sense and respond to a variety of biotic and abiotic stresses. The architecture of root systems and their growth are known to be strongly affected by the environmental conditions found in the soil. However, the acquisition of cell identities at the root meristem is still mainly viewed as ontogenetically driven, where a small number of stem cells generate all the cell types through stereotyped divisions followed by differentiation, along a simple developmental trajectory. The extent to which environmental cues precisely shape and affect these developmental trajectories remains an open question. We used single-cell RNA-seq, combined with spatial mapping, to deeply explore the trajectories of cell states at the tip of *Arabidopsis* roots, known to contain multiple developing lineages. Surprisingly, we found that most lineage trajectories exhibit a stereotyped bifid topology with two developmental trajectories rather than one. The formation of one of the trajectories is driven by a strong and specific activation of genes involved in the responses to various environmental stimuli, that affects only of a subset of the cells in multiple cell types simultaneously, revealing another layer of patterning of cell identities in the root that is independent of cell ontogeny. We demonstrate the robustness of this environmentally responsive transcriptional state by showing that it is present in a mutant where cell type identities are greatly perturbed, as well as in different *Arabidopsis* ecotypes. We also show that the root can adapt the proportion of cells that acquire this particular state in response to environmental signals such as nutrient availability. The discovery of this cell state reveals new layers of cell identity that may underpin the adaptive potential of plant development.

## Main text

The root of *Arabidopsis thaliana* is a powerful model for studying cell differentiation in plants. In this organ, cell types are arranged in concentric layers around a central vascular bundle with a bilateral symmetry (Fig. 1A). Numerous morphological and genetic studies have demonstrated that each cell type corresponds to a cell lineage that is generated by a specific stem cell population maintained near the tip, in a region called the meristem ^1^. Within the cell lineages, with each new stem cell division, cells are displaced upwards and differentiate, thereby creating a longitudinal developmental trajectory where the spatial arrangement of cells recapitulates the temporal dynamics of differentiation (Fig. 1A).

**Fig. 1.**
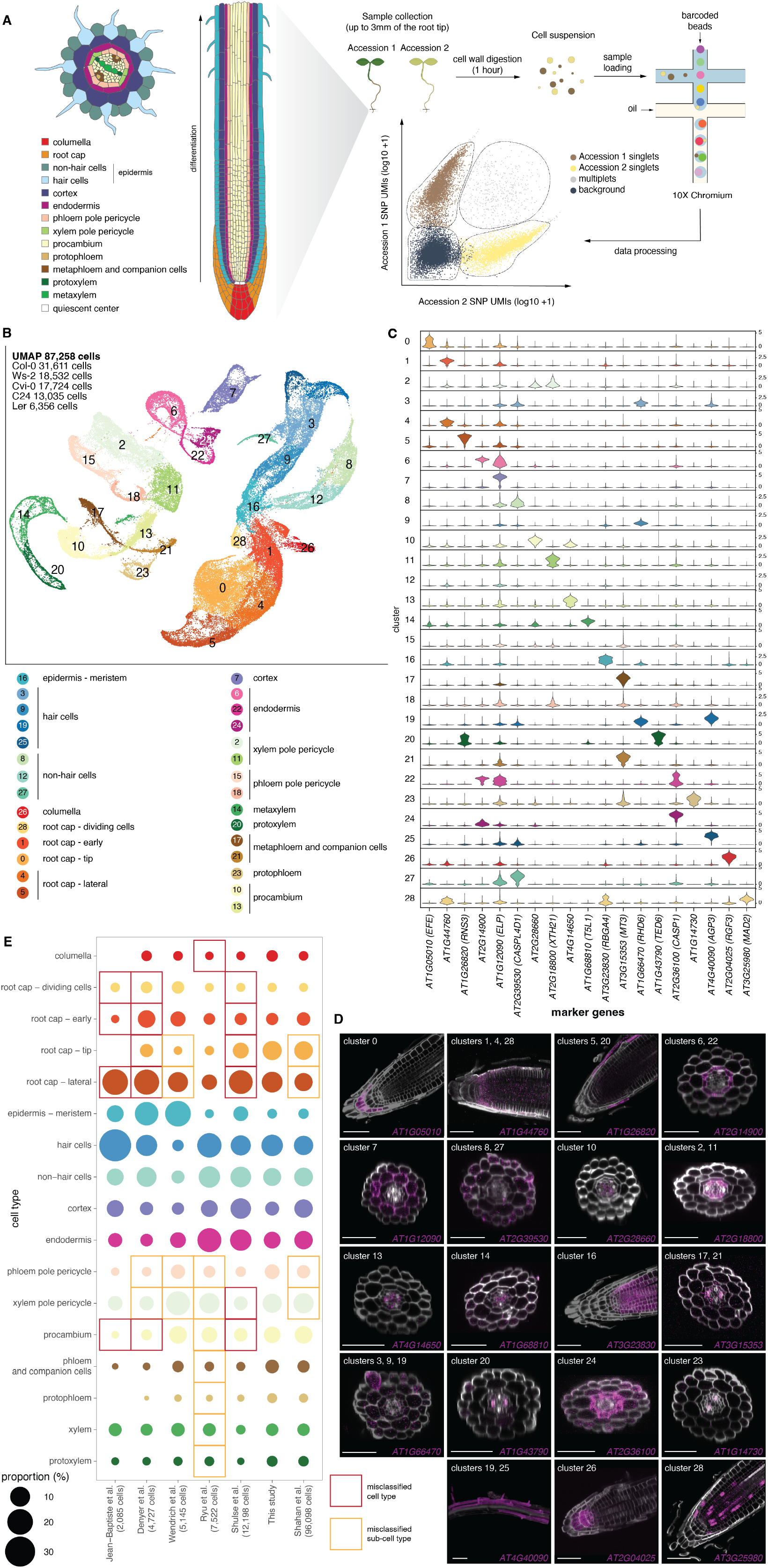
A high-resolution single-cell atlas of the *Arabidopsis* root tip. (**A**) Schematic representation of the root tip, the experimental procedures to obtain protoplasts and perform single-cell RNA-seq, and *in silico* strategy for identifying singlets and multiplets. (**B**) UMAP plot of our reference atlas of 87,258 cells from five *Arabidopsis* accessions. Cells are colored by clusters whose identity is determined by whole-mount in situ hybridization chain reaction (HCR). (**C**) Violin plot showing the cluster-specific expression of 19 markers used for determining cluster identity via whole-mount HCR. (**D**) Whole-mount HCR of the 19 markers identified in (C) for precise cell type assignment of the clusters. Scale bar = 50µm. (**E**) Dotplot showing the representation in published root single-cell datasets of the different root cell types we identified in this study. Dot size represents the percentage of cells within a cell type. Cell assignments were obtained by transferring cell labels from our reference atlas. Red rectangles indicate cell types detected in our atlas but not detected in the published study’s original assignment, and orange rectangles show cell types that were detected in the other study but at a lower definition of cell identity.

While transcriptional characterisation of root cell populations has previously focused on the analysis of bulk sorted lineages, the recent development of single-cell technologies is enabling more precise investigation of the spatio-temporal diversity of cellular states ^2–6^. Single-cell RNA-sequencing (scRNA-seq) has recently been used to better define the gene expression signatures of the known cell types of the *Arabidopsis* root and, by arranging cells of each lineage in developmental trajectories, more finely define the dynamics of gene expression during cell differentiation ^7–14^. While this has been crucial for identifying transient cellular states in some cell lineages, these studies have treated each lineage independently, overlooking potential layers of gene regulation that do not depend on the developmental origin of the cell.

Here, we map each cellular state of the root, providing the most precise root cell atlas to date, by a combination of scRNA-seq and whole-mount *in situ* hybridization chain reaction (HCR) ^15^. We show that many cell lineages exhibit a more complex topology than anticipated by using cell trajectory analyses. We determine that these topologies are a result of a strong transcriptional signature superimposed upon the developmental identity of a subset of the cells in each lineage, and controlled by environmental cues and plant hormones. As a result, cell types in the root are in fact composed of subsets of cells whose function is modulated in response to environmental information perceived by the plant.

### Results

### A high-resolution atlas of *Arabidopsis* root cell lineages reveals uncharacterized branching events in developmental trajectories

To characterize cell states and their dynamics during root development, we generated a high-resolution scRNA-seq (10x Genomics) atlas of the wild type *Arabidopsis* root tip. Protoplasts were isolated from root tips of two different *Arabidopsis* ecotypes, mixed at the same concentration, and processed for single cell transcriptome profiling to enable disambiguation of *bona fide* single cells from background and doublets (see Methods) (Fig. 1A, fig. S1, A to E). We obtained a total of 87,258 wild-type cells from 5 different ecotypes (accessions), and detected a total of 25,738 genes, with a median of 10,255 unique molecular identifiers (UMIs) and 3,199 genes detected per cell. After excluding 1,859 genes we detected as significantly affected by protoplast isolation (see Methods), we removed batch- and genotype-specific differences to create a reference atlas (Table S1, Fig. 1B) ^16^.

We performed clustering and differential expression analyses to characterize the cellular heterogeneity within our dataset. In a first attempt to assign an identity to the 29 resulting clusters, we compared our cluster-specific genes with published gene expression profiles of sorted populations of root cell types expressing fluorescent reporters (fig. S2) ^6^. Although some of the markers from sorted cells intersected with our cluster markers and allowed potential identification of some cell types, the dataset did not provide any specific marker for the lateral root cap, procambium, and xylem pole pericycle clusters. Moreover, the markers of several of our clusters were not enriched specifically in any sorted populations, making it difficult to clearly assign an identity. To circumvent this difficulty, we adapted a whole-mount HCR ^15^ method to optimize it for *Arabidopsis* root tips and combined it with a clearing technique ^17^ in order to precisely map the expression of the 19 most specific cluster markers of our dataset (Fig. 1C). Using this method, we could identify all the known cell types of the root and, beyond this, subpopulations of the root cap (early, tip, and lateral) (Fig. 1D). Only 2 genes (*AT3G23830* and *AT3G25980*) did not have cell type-specific expression patterns when examined by HCR, contrasting with their definition as cluster-specific markers. However, their associated clusters could be identified as the meristematic portion of the epidermis (cluster 16) and early dividing cells in root cap formation (cluster 28) (see Methods). We annotated these clusters as “epidermis - meristem” and “root cap - dividing cells”.

To demonstrate the value of our reference atlas and the precision of cell identification obtained with the *in situ* hybridization, we used it to annotate cells from all published *Arabidopsis* root tip datasets ^7–10,12,13^, with the Seurat label transfer method (Fig. 1E), and compared it to their original assignment. While all the cell types we characterized are present in the published studies except one, their original assignment was less precise. All other studies were missing at least one cell type or sub-cell type that we identified, such as the pericycle subdivision into phloem pole pericycle (PPP) and xylem pole pericycle (XPP), potentially due to their reliance on bulk sorted population gene expression data to assign cell identities, which we determined does not allow identification of all cell types of the root. This demonstrates the importance of precisely spatially resolving our single-cell data to validate cell states independently.

A striking feature of our dataset (Fig. 1B) is the clear grouping of cells according to their developmental lineages (epidermis - root cap, cortex, endodermis, XPP, PPP, procambium, phloem, and xylem), confirming that the ontogenic signal is the strongest transcriptional component of cell identity. Within each of the lineages, cells can be ordered in trajectories that recapitulate the developmental transitions from an undifferentiated to a differentiated state. For this trajectory analysis, we focused on the Col-0 accession cells only, and subsetted the dataset into eight lineages that were treated independently (Fig. 2A). For each lineage, we performed sub-clustering of the cells (Fig. 2A) and calculated a “stemness score” (see Methods) that allowed identification of the cell with the most meristematic state, which was then considered as the starting point (t=0) of the trajectory (Fig. 2B, fig. S4). A partition-based graph abstraction (PAGA) method ^18^ was used to infer putative transitions between clusters, thereby identifying the topology of the cell lineages (Fig. 2C), and the developmental directionality of the trajectories was determined by computing the diffusion pseudotime ^18–20^ from t=0 (Fig. 2B). The combination of both the PAGA and pseudotime reveals all the potential branching events occurring during development (Fig. 2C).

**Fig. 2.**
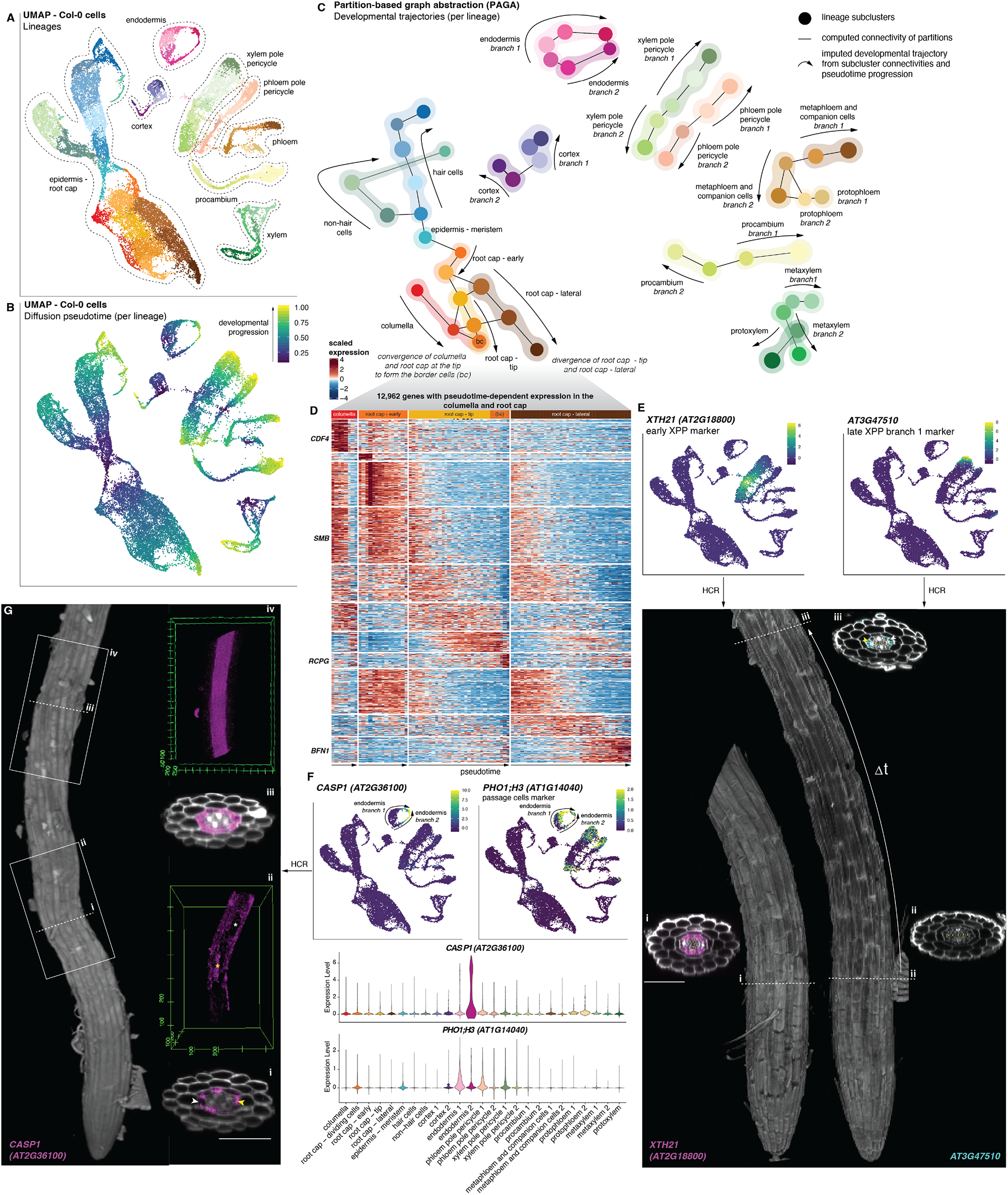
Pseudotime analysis reveals unexpected complexity in cell lineage trajectories. (**A**) UMAP plot of the Col-0 cells showing the different cell lineage subdivisions treated independently in the pseudotime analyses. Cells are colored by subclusters in each lineage. (**B**) UMAP plot showing the diffusion pseudotime assignment for each cell, per lineage. t=0 corresponds to the cell with the highest “stemness score” (see fig. S5) and t=1 represents the most distinct state from t=0 in the lineage. (**C**) Developmental trajectories obtained by combining the connectivity between the subclusters shown in (A) computed in a partition based graph abstraction (PAGA) and the pseudotime inference calculated in (B) reveal two distinct developmental branches affecting most of the cell lineages (named branch 1 and 2). (**D**) Pseudotime-dependent expression in the columella and root cap. Heatmap showing clusters of branch- and pseudotime-specific genes affecting the four different branches of the root cap. Examples of genes known to be expressed at specific stages are labeled on the left. Cells were ordered in pseudotime for each branch and binned in bins of 100-cells. (**E**) UMAP plots (top) and confocal images (bottom) of whole-mount HCR showing the expression of two XPP markers: *XTH21* (left) and *AT3G47510* (right). Optical transverse sections (i for *XTH21*; ii and iii for *AT3G47510*) show that *AT3G47510* is detected (iii) at much later developmental stages (Lit) than *XTH21* (i). While *XTH21* is detected homogeneously in the XPP (i), *AT3G47510* is only detected in a portion of the XPP cells (iii, yellow arrow), with some cells having no fluorescent signal (iii, white arrow). Yellow “x” marks xylem cells. Scale bar = 50µm. (**F**) UMAPs and violin plots showing the expression of *CASP1* and *PH01;H3* in the endodermis. *CASP1* is enriched in the developmental branch 2, and *PH01;H3* in branch 1. (**G**) 3D views and optical sections showing the expression of *CASP1* at the root tip by whole-mount HCR. *CASP1* is expressed only in a subset of the endodermal cells close to the meristem (i and ii), and homogeneously in the endodermis at a more differentiated stage (iii and iv). Yellow and white arrows indicate positive and negative cells respectively. Scale bar = 50µm.

The “epidermis - root cap” lineage exhibits the most complex topology, with the inferred trajectories recapitulating known developmental events: the split of epidermal cells into hair cells and non-hair cells, and the split of the root cap into a part that will grow laterally and a part that will grow towards the tip and converge with columella cells to form the sloughing border cells ^21^ (Fig. 2C). While those branching events are known to happen *in vivo*, to date no root single-cell analysis has been able to identify them, further highlighting the resolution of our cell annotations and trajectory analyses. We used the root cap trajectories to demonstrate that we can identify trajectory- and pseudotime-specific variable genes, including genes known to be involved at different stages of root cap development, for instance the Dof transcription factor (TF) *CDF4* expressed in the differentiating columella ^22^, *SMB* activated in early root cap development ^23^, the *ROOT CAP POLYGALACTURONASE* (*RCPG*) expressed at the tip, or *BFN1* expressed laterally at the programmed cell death site II ^24^, further validating the imputed developmental trajectories (Fig. 2D).

For all the other lineages, except phloem and xylem that are composed of different cell types, the trajectories are expected to be linear with a single transition from an undifferentiated to a differentiated state, recapitulating the longitudinal arrangement of cells. Unexpectedly, the PAGA analysis shows that all other cell types of the root have not one but two branches in their trajectory that diverge at a very early pseudotime point (Fig. 2, B and C). To independently verify that this topology really corresponds to two branches (“branch 1” and “branch 2”) diverging from an earlier developmental starting point, and not a misassignment of t=0, we performed whole-mount HCR of two genes that are expressed in early and late stages of XPP development based on the scRNA-seq data (Fig. 2E). We observe that *XTH21*, which is expected to be expressed early in both XPP branches, is indeed detected in all XPP cells close to the meristem, while the late XPP branch 1 marker *AT3G47510* only starts to be detected much further away from the root tip and is not homogeneously detected across XPP cells. This confirms the branching nature of these trajectories over development, which reveals two previously unidentified *in vivo* alternate cellular states.

We also validated the endodermis trajectory by looking at the expression pattern of the casparian strip formation regulator *CASP1* by whole-mount HCR (Fig. 2F). *CASP1* is strongly expressed at a late converging state between branch 1 and branch 2 in the endodermis, but starts to be expressed earlier in branch 2 endodermis cells. We found that *CASP1* mRNA is indeed strongly detected throughout the endodermis in late differentiated states, but was found to be detected in only a subset of the cells closer to the meristem (Fig. 2G). While *CASP* genes tend to be enriched in branch 2, branch 1 is enriched for *PHO1;H3*, a marker for passage cells ^25^, which are a specific population of un-suberized cells in the differentiated endodermis that act as cellular gatekeeper, controlling access to the root interior ^26^. Thus, the branching event we observe early in endodermis formation might be involved in the specification of passage cells before the onset of suberisation.

Importantly, when we assign identities to cells from published *Arabidopsis* root scRNA-seq datasets using our reference atlas and incorporating the branch identities, all datasets have cells assigned to both branches for all affected cell types, with similar genes found to be branch-specific (Fig. S5), demonstrating that this diversity of cell states is a robust feature of roots when characterized at single cell resolution.

While we expected to observe gradients of gene expression in each root lineage that recapitulate changing cellular states throughout differentiation, the observation of two alternate states on top of the pseudotime differentiation axes of multiple lineages demonstrates that another level of complexity exists in most cell types, which was not previously anticipated.

### A common transcriptional signature drives the branching of multiple lineages

In order to globally identify the cell type-, branch-, and pseudotime-specific transcriptional signatures, and the specificities and commonalities between them at the whole root level, we used self-organizing maps (SOM) to detect modules of co-expressed genes in our Col-0 dataset. We identified 64 distinct modules, composed of 78 to 791 genes each (Table S2), and visualized their average scaled expression across the eight cell lineages over pseudotime (Fig. 3A) and their associated GO terms (Fig. 3B).

**Fig. 3.**
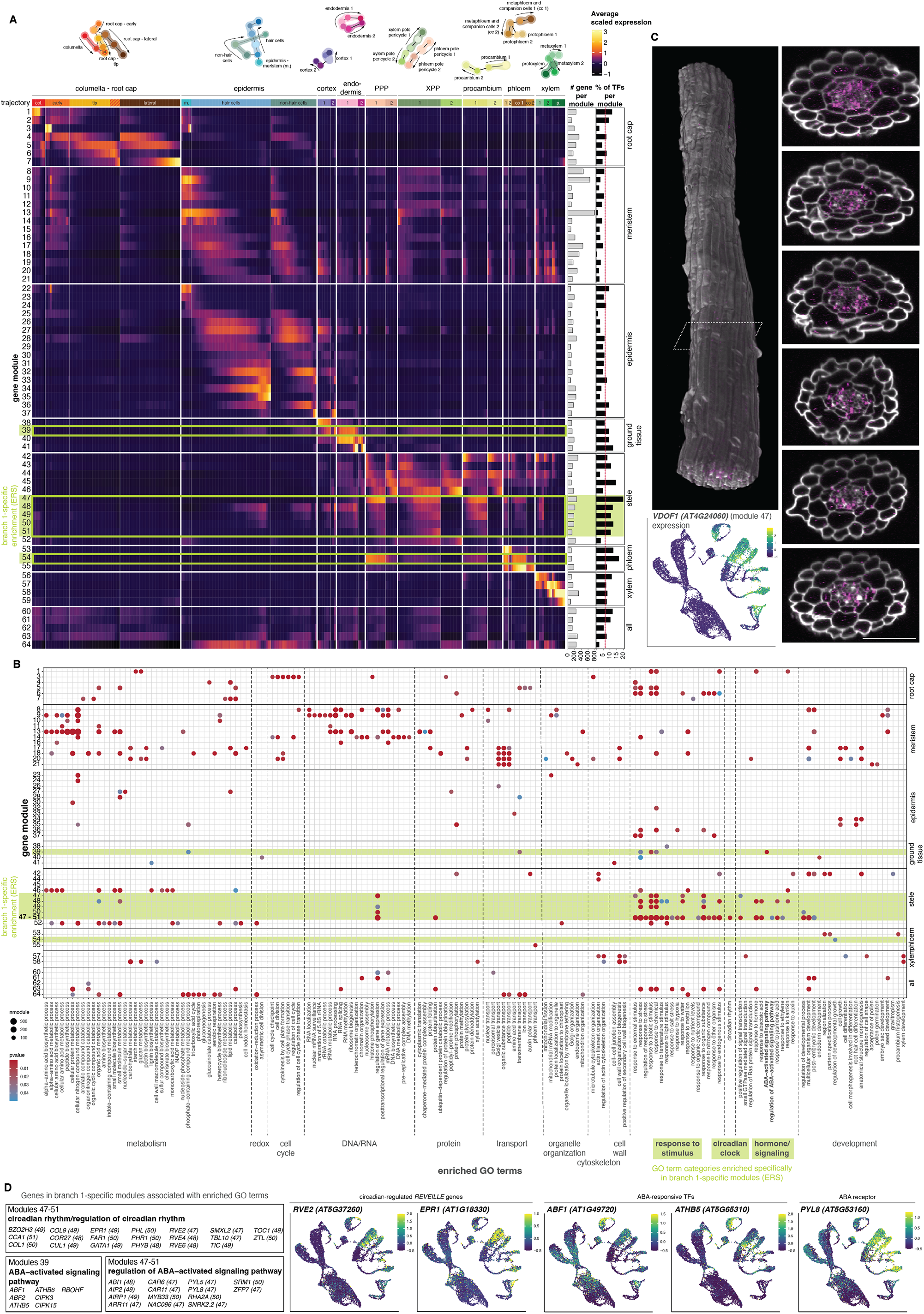
Co-expression gene modules identify a branch-specific transcriptional signature common to multiple cell types. (**A**) Heatmap showing the expression pattern of the 64 modules of co-expressed genes obtained from the self-organizing map (SOM) analysis. Cells were ordered in pseudotime, per developmental branch, and grouped in bins of 100-cells each. Barplots show the number of genes and transcription factors (TFs) per module. The cell types the modules are enriched in are listed on the right. Branch 1-specific modules are highlighted in green. (**B**) Dotplot of enriched GO terms per co-expression gene module described in (A). Terms associated with the branch-1 specific (environmentally responsive state [ERS]) modules are highlighted in green. (**C**) UMAP plot and confocal images showing the expression of *VDOF1*, which belongs to a branch-1 specific module, in the scRNA-seq dataset and *in vivo* detection and visualization by whole mount HCR. Multiple transverse optical sections along the longitudinal axis of the root are shown. Scale bar = 50µm. (**D**) Focus on the genes associated with the ABA- and circadian-related terms, which are specifically enriched in branch-1 specific modules.

Modules specific to each cell type of the root were detected, many of which contain genes known to be expressed in the respective lineages. For instance, genes related to secondary cell wall formation and xylem development were enriched in xylem specific modules (57, 58), the “root hair differentiation” term was enriched in epidermal modules 34-35, and the casparian strip formation genes *CASP1-5* associated with the “cell-cell junction assembly” term were enriched in the late converging endodermis state and earlier in branch 2 (module 41) (Fig. 3B, Table S2).

We found branch-specific gene modules for all lineages that exhibited a branch split, from the endodermis inwards. For instance, module 39 is composed of genes particularly enriched in branch 1 of the endodermis but not branch 2, and module 54 is enriched in branch 1 of phloem and PPP lineages (Fig. 3A). Crucially, we found that the majority of the branch-specific modules affect multiple cell types simultaneously, and are always enriched in the same branch. We identified a group of five modules (modules 47-51) that are specifically enriched in branch 1 of multiple cell types (endodermis, PPP, XPP, procambium, phloem, and xylem), revealing a common gene expression signature specific to a subset of the cells in all of these distinct lineages, regardless of their developmental origin (Fig. 3A, fig. S6), and which would be responsible for the bifurcation observed in the PAGA trajectories (Fig. 2C). We note that, although the branch 1-specific genes are always enriched in multiple lineages simultaneously, some variation can be observed in the level of enrichment across different cell types, suggesting that this transcriptional state is not exactly equivalent in all cell types. This hypothesis of a common co-expression signature superimposed upon distinct developmental identities is further supported by the observation that the number of genes and transcripts detected in branch 1 as compared to branch 2 is systematically higher, in all cell types (fig. S7, A and B). However, the formation of the branches is not a result of a coverage (number of UMIs per cell) artefact, as demonstrated by downsampling analyses (fig. S8). This is also observed for cells assigned as branch 1 or branch 2 in our reanalysis of published *Arabidopsis* root scRNA-seq datasets (fig. S7C), although the branching and associated transcriptional states were not previously discerned in these studies. Moreover, the five branch-specific modules, and in particular module 47, contain a high proportion of transcription factors, which could underpin the increase in detected transcripts and genes specifically in those cells (Fig. 3A).

We note that although the PAGA analysis did not detect two branches for the epidermal hair cell and non-hair cell trajectories, a small branch with the lowest number of genes and UMIs that does not express the branch 1-specific genes can also be observed in the epidermal cell types, suggesting that higher resolution profiling might detect analogous epidermal branching (fig. S8, A and B). In the cortex, however, the branching event identified in the pseudotime analysis does not appear to result from the same superimposed state that is common to other lineages, and was not considered in subsequent analyses.

Assessment of protoplast isolation responsive-genes verified that this branch-specific gene signature is not due to sample preparation, since genes in these modules were not particularly up- or down-regulated after cell dissociation (fig. S9). Importantly, we independently validated the existence of this cellular state *in vivo* in a subset of the cells of many distinct types by performing whole-mount HCR of one of the branch 1-specific genes of module 47, the Dof transcription factor *VDOF1* (Fig. 3C). We confirmed that *VDOF1* is detected in multiple cell types simultaneously, and only in a subset of the cells. Notably, we did not detect a specific spatial arrangement of cells that express the gene, with the radial expression exhibiting variability along the longitudinal axis.

Together, these co-expressed gene modules demonstrate that a gene co-expression signature common to a subset of cells from multiple lineages drives the formation of the branch 1 state, highlighting another layer of regulation of cell identities at the root tip, independent from the developmental lineages.

### The branch-specific transcriptional signature is linked to environmental responses

To investigate the function and origin of this gene co-expression signature we assessed gene ontology (GO) terms enriched in the different branch-specific modules, including by grouping the branch-specific modules 47-51 that display highly similar enrichment patterns (Fig. 3C). Branch 1-specific genes are strongly associated with processes involved in response to stimuli and response to hormones, in particular to abscisic acid (ABA). Module 39, specifically enriched in the endodermal branch 1, contains 4 ABA-responsive TFs (ABF1, ABF2, ATHB-5, and ATHB-6) known to mediate transcriptional regulation of ABA stress responses ^27–31^, suggesting that this hormone pathway is particularly active in this subset of the endodermis cells. In modules 47-51, which are specifically enriched in branch 1 in multiple stele cell types, 50% of the genes associated with the term “response to hormone” are linked to other GO terms involving ABA (fig. S10). In particular, 14 genes involved in the “regulation of abscisic acid-activated signaling pathway” were found in these modules, including two members the C2-DOMAIN ABA-RELATED protein family (*CAR6, CAR11*), known to enhance ABA sensitivity by targeting its receptors to the plasma membrane ^32^, two ABA receptors (*PYL5, PYL8*) ^33,34^, the co-receptor and negative regulator *ABI1* ^*35–37*^, the kinase *SRK2D* ^*38,39*^, and three transcription factors known to mediate some ABA responses (*MYB33, NAC096, SRM1*) ^40–42^ (Fig. 3D, fig. S11). The receptor PYL8 seems to be particularly important for the regulation of ABA signaling in the root, compared to the other member of the PYL multigenic family. PYL8 levels are specifically increased by ABA treatments while no significant effect was observed for PYL1 and PYL4 ^43^, and PYL8 is the only receptor of this family that known to play a non-redundant role in ABA sensitivity in the root ^44^. Altogether, this suggests that ABA signaling pathways are more active in the branch 1 cells and may be driving, at least partially, their specific patterns of gene expression.

In addition to these hormone response links, the GO terms “circadian clock” and “regulation of circadian clock” were found exclusively enriched in the branch 1-specific modules (Fig. 3C). Among the branch 1 module expressed genes associated with these terms, we found the core clock genes *TOC1* and *CCA1* ^*45*^, as well as other modulators of the clock including *ZTL* ^*46*^, *BZO2H3* ^*47*^, the effectors of light signaling and entrainment of the clock *FAR1* ^*48*^ and *PHYB* ^*49*^, and the members of the REVEILLE family *RVE2, RVE4, RVE6*, and *EPR1* ^*50– 52*^, the expression of which is clock-regulated (Fig. 3D, fig. S11). This strongly indicates a potential circadian input or output to the branch-specific state.

Overall these functional enrichment analyses strongly suggest that the cellular state responsible for the branching events of multiple lineages is linked to responses to various environmental stimuli, therefore henceforth it will be referred to as the “environmentally responsive state” (ERS), with “branch 1” and “branch 2” corresponding to ERS-positive (ERS+) and ERS-negative (ERS-) cells, respectively.

To investigate what is driving the ERS, we sought to identify putative cell-autonomous regulators of the different gene modules, by selecting TFs whose predicted binding target genes are enriched in the same module the TF is expressed in, using the *Arabidopsis* DAP-seq dataset ^53^ and the predicted binding similarities from Cis-BP2 ^54^. We identified 104 putative transcriptional regulators for 31 of the gene modules (Fig. 4A). Among these predicted cell-autonomous regulators, 42 were previously described as playing a role in the development of the cell type they are enriched in, indicating our approach is effective at identifying important regulators.

**Fig. 4.**
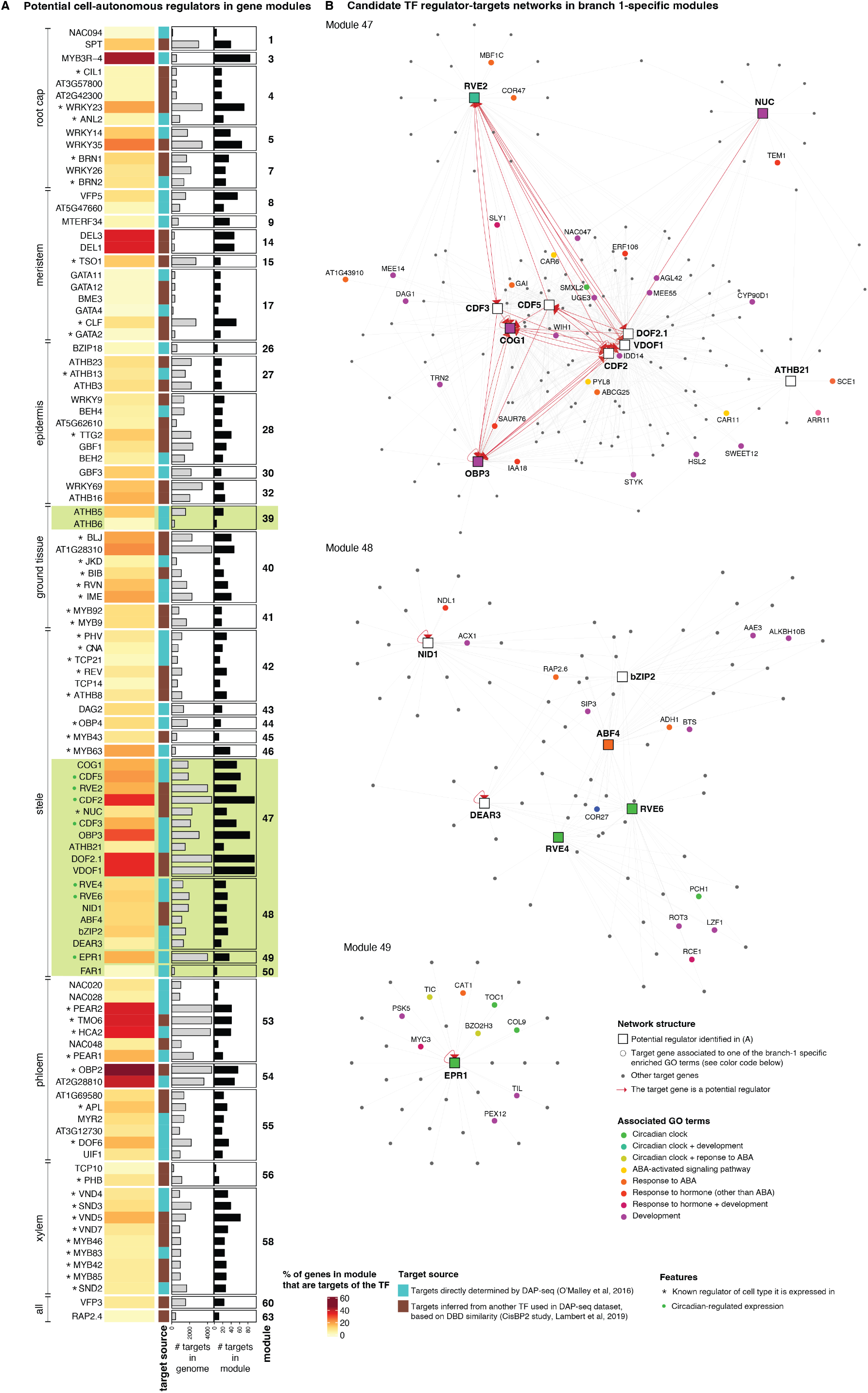
Putative cell-autonomous regulators of the ERS confirm ASA-responsive and circadian signature. (**A**)Predicted cell-autonomous regulators (TFs) of the co-expression gene modules, identified by enrichment of TF binding target genes in the modules (numbered on left) that they are expressed in (Fisher’s exact test, FDR <0.05). Heatmap shows the proportion of genes in each module that are identified as targets of each listed TF regulator, and barplots show the number of predicted binding targets in the genome and in the module for each TF. The target source color key indicates whether the targets have been determined directly for each TF by DAP-seq or imputed from a similar TF using the cis-BP binding similarity predictions. Asterisks and circles indicate known developmental regulators of the cell type the TF is enriched in and circadian-regulated genes, respectively. (**B**) TF-target networks in ERS modules 47, 48, and 49. Putative cell-autonomous regulators (squares) are linked to their predicted targets (circle) by an arrow. Regulators that are also targets are indicated by red arrows. Only genes associated with the enriched ABA-, hormone-, development- and circadian-related terms are highlighted.

Candidate regulators identified for the branch-specific modules are in agreement with the enriched GO terms identified above (Fig. 4A). The ABA-responsive ATHB5 and ATHB6 were found in the endodermis specific module 39. In the modules 47-51 enriched in the ERS+ cells of multiple cell types, we identified several TFs whose expression is circadian-regulated, including a group of Dof cycling factors *CDF2, CDF3*, and *CDF5*, and the REVEILLE genes *RVE2, RVE4, RVE6*, and *EPR1* that encode known modulators of the clock. We also found the mediator of ABA responses *ABF4*. More generally the ERS-specific modules 47-51 are enriched for genes with an ABA-responsive element binding motif in their vicinity (Table S3). This TF-binding analysis further demonstrates the involvement of ABA and the circadian clock in the regulation of this ERS, as genes enriched specifically in this cellular state are not only annotated as part of their biological processes but also putative targets of their regulators based on TF DNA binding data.

Next, to better predict the effective cell-autonomous TF targets *in vivo* and the structure of the regulatory network per module, we combined the genome-wide binding sites predicted by DAP-seq and the co-expression modules identified by scRNA-seq. We investigated the TF-target gene network of the ERS-specific modules 47-49, revealing links between the module genes belonging to the enriched ABA, hormonal, developmental, and circadian response GO terms (Fig. 4B). Although the circadian clock has been shown to regulate ABA biosynthesis and signaling, and ABA has been demonstrated to feedback on the clock ^55^, we only found a putative link between this pathways at the root tip: several cycling Dof and also RVE2 are predicted as regulating the expression of *CAR6, CAR11* and *PYL8*, which are three enhancers of ABA sensitivity.

In sum, the ERS that is present in only a proportion of cells regardless of their lineage is strongly linked to pathways involved in the plant response to its environment, in particular through a specific signature of activation of ABA-related genes, a hormone known to play a crucial role in the response to stresses, as well as, unexpectedly, various key regulators of circadian responses.

### Branching events are maintained despite developmental perturbations

We hypothesized that the ERS is superimposed on, and independent from, developmental identities because of the observation that it affects multiple cell types. Therefore, we next asked whether the ERS would be maintained upon developmental perturbations of cell types. To do so, we investigated cell identity alterations in the *scr-3* line that has a premature stop codon in the *SCARECROW (SCR)* gene ^56^, which encodes a transcription factor that controls various cell identity processes in the endodermis and stele ^1^, which exhibit the ERS.

We first characterized global changes in cell type composition by scRNA-seq of the *scr-3* mutant root tip (Fig. 5A), using the Col-0 cells of our reference atlas to assign identities to the mutant cells. Multiple cell types are affected in the *scr* mutant (Fig. 5B, fig. S12). As expected, both the endodermis and protoxylem were strongly depleted in *scr-3* as compared to our five Col-0 replicates, and correspond to two well-characterized phenotypes of *scr*: the presence of a single ground tissue layer ^57,58^ and the ectopic formation of metaxylem in place of protoxylem ^59^. We also found other unexpected cell types to be affected in the mutant (Fig. 5B, fig. S12). First, the columella, which was not reported to be altered in a previous study characterizing the single-cell phenotype of *scr-4* ^*13*^, is significantly depleted in *scr-3*. This is consistent with a previous report showing that columella stem cells in *scr-1* abnormally express markers of differentiated root cap cells ^60^. Second, both the XPP and PPP identities are strongly reduced in the mutant (Fig. 5B). While the columella and pericycle cell types are depleted in the *scr-3* dataset, their corresponding layers are present physically in *scr-3* ^57^. Thus, this phenotype is not caused by a morphological defect, but rather by the modification of the cell identities. We found that the root cap and procambium cells, which are cell types adjacent to the columella and pericycle in the root, are significantly over-represented in the mutant (Fig. 5B, fig. S12). We hypothesize that the cells that physically correspond to columella and pericycle in the wild-type root have respectively acquired a root cap and procambium identity in *scr-3*. Therefore, we demonstrate that a mutation in the *SCR* gene leads to a broad range of cell identity alterations, some of which were not previously described. Despite these major alterations in cell identity, with some cell types lost and others present ectopically compared to wild type, we detect cells assigned to the ERS for all cell types of the stele (Fig. 5C) and observe its associated branching in the UMAP representation.

**Fig. 5.**
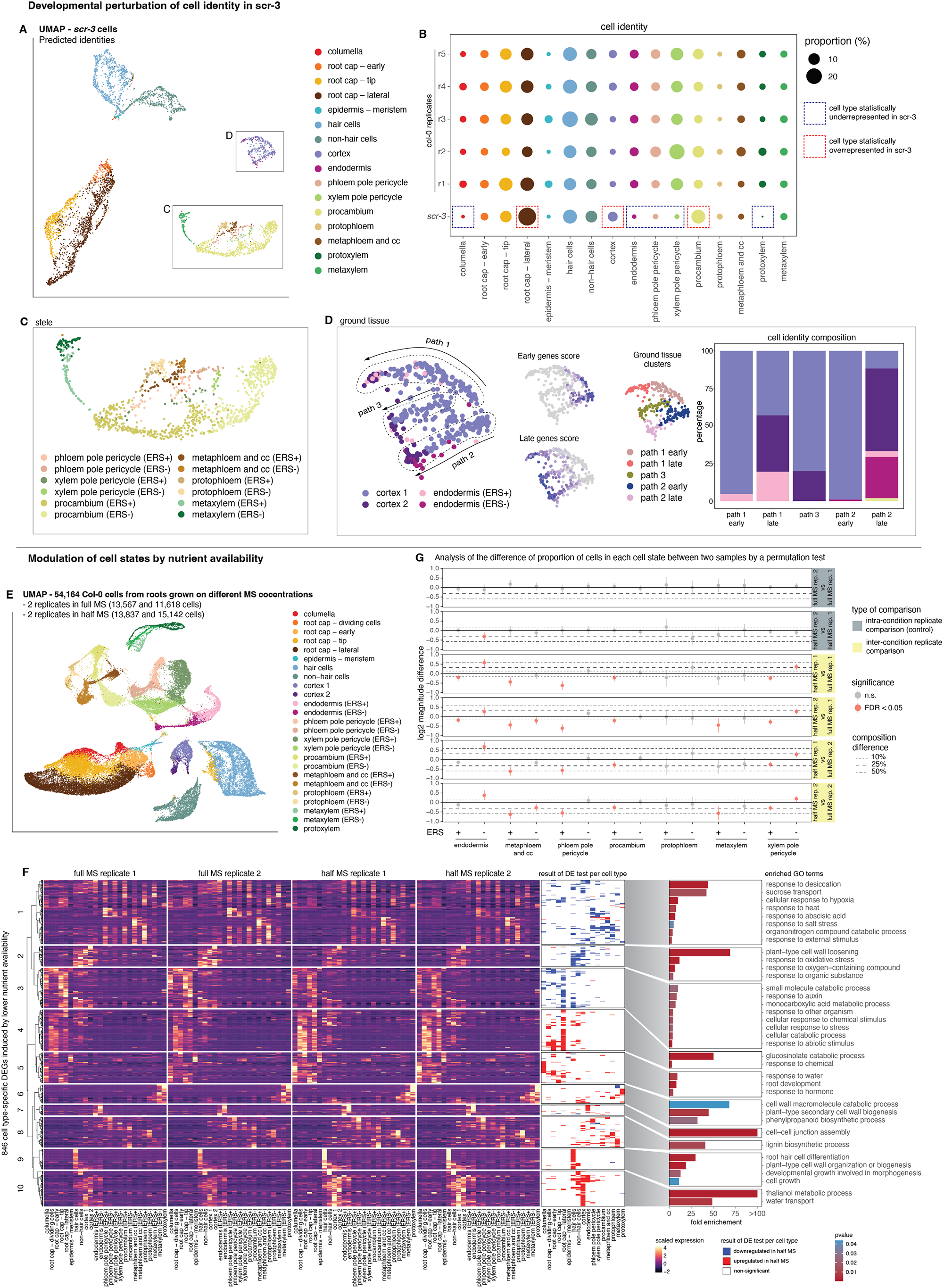
The ERS is maintained upon developmental perturbations of cell identities and modulated by nutrient availability. (**A**) UMAP plot of cells from *scr-3* root tip scRNA-seq. Cells are colored by the cell type they have been assigned to, using our Col-0 reference atlas. (**B**) Dotplot showing the proportion of cells of each cell type in the mutant as compared to the five Col-0 replicates. Cell types significantly over- or under-represented (permutation test, FDR<0.05) in *scr-3* as compared to each of the Col-0 replicates are indicated (see fig. S12). (**C**) Close-up of the stele cells in the *scr-3* UMAP plot (indicated by box in panel (A)). Cell assignment takes into account the environmentally responsive state (ERS) branching events. (**D**) Close-up of the ground tissue cells in the *scr-3* UMAP plot (indicated by box in panel (A)). An “early” and “late” gene score (similar to that shown in fig. S5) has been calculated to infer trajectory direction in *scr-3* ground tissue cells. Three developmental paths are detected. Ground tissue cells are grouped in five different clusters (path 1 early, path 1 late, path 2 early, path 2 late, path 3). Stacked barplot shows the proportion of cells in each path that has been assigned to the cortex 1, cortex 2, endodermis 1, and endodermis 2 identities. (**E**) UMAP plot of cells from root tips grown in different nutrient availability conditions: full (1x) or half (0.5x) concentration Murashige and Skoog (MS) media (2 replicates per condition). Cells are colored by cell state. (**F**) Comparison of cell state (ERS+ or ERS-) proportion between replicates of the same MS concentration (control) or different MS concentration conditions. Point-range plot showing the confidence interval for the cell state (ERS+ or ERS-) proportional difference between replicates, as compared to 1000 random permutations, to account for subsampling effects (see Methods). Significantly different proportions (permutation test, FDR<0.05) are indicated in pink. (**G**) Heatmap showing the scaled average expression per cell state (ERS+ or ERS-) of genes differentially expressed (DE) between the half (0.5x) and full (1x) MS media conditions, in at least one cell type. Genes are clustered by their expression pattern. Barplots indicate enriched GO terms per cluster, colored by enrichment p-value.

In the mutant ground tissue layer, we observe that cells transition from a cortex to an endodermis identity, as recently reported by scRNA-seq characterization ^13^. Despite the endodermis identity not being present at early stages of development, both ERS+ and ERS-endodermal cells are detected in our *scr-3* dataset (Fig. 5D). Three different developmental paths for the ground tissue layer are apparent: one that will differentiate into ERS+ endodermis, another into ERS-endodermis, and a third that will remain cortex (Fig. 5D). This further supports that the branching events caused by the ERS are not regulated developmentally and formed by division events early in development, but rather are states superimposed on multiple cell lineage identities regardless of their cell type and developmental stage.

### Nutrient availability modulates the proportion of ERS+ cells

The genes and GO terms enriched in the ERS+ cells are strongly associated with responses to various environmental stimuli, and because of the link between the branching events and the passage cell marker *PHO1;H3* (Fig. 2F), whose expression is affected under phosphate-, zinc-, and iron-deficiency conditions ^25,61^, we next aimed to characterize the effect of nutrient availability on the ERS.

We utilized scRNA-seq to compare the composition of cell transcriptional states in root tips grown on two different concentrations of plant basal growth media (full [1x] versus half [0.5x] concentration Murashige and Skoog [MS] media), thereby subjecting the plants to different concentrations of nutrients and vitamins. This range is commonly used in *Arabidopsis* studies and does not drastically affect root growth. We generated a dataset of 54,164 single cells, with 25,185 from seedlings grown on full MS and 28,979 from seedlings on half MS, each condition having two biological replicates (Fig. 5E). All cell types were detected in the dataset, at a similar proportion for each condition and replicate (fig. S13, A and B). We investigated differentially expressed genes (DEGs) between the two nutrient conditions for each cell type. Lower nutrient availability induced cell-type specific changes in the expression of 846 genes (average log fold-change ≥ 0.25, adjusted p-value < 0.05, t-test) (Fig. 5F, Table S4). For example, we detected an increase in the expression of genes involved in trichoblast formation in hair cells (cluster 9), multiple *CASP* genes in the endodermis (cluster 7), lignin biosynthetic regulators in multiple cell types (cluster 8), and various water transporters enriched in the cortex (cluster 10). The modulation of the formation of hair cells and the casparian strip and the ability to transport water are expected to be affected by changes in nutrient availability, and seedlings grown on half MS had a higher density of hair cells (fig. S14). We found that 174 (20%) of the DEGs are specifically enriched in ERS+ cells, down-regulated under lower nutrient availability conditions (cluster 1), and associated with GO terms linked to response to environmental stimuli and ABA (Fig. 5F). This suggests that the proportion of cells affected by this superimposed state could be modulated under different nutrient availability concentrations. To test this hypothesis we compared the proportion of cells assigned as ERS+ vs ERS-for the endodermis and stele cell types, in the two different nutrient concentrations. We used a permutation test to compare each replicate of the dataset, allowing us to account for any sub-sampling effect that could contribute to the differences in proportion of cells in each branch (Fig. 5G). We found that ERS+ cells are significantly depleted (FDR < 0.05) in half MS as compared to full MS conditions in most cell types tested, consistent with the down-regulation of ERS-specific genes (Fig. 5F). We also observed a significant increase (FDR < 0.05) in the proportion of cells assigned as ERS-endodermis cells, which expresses the casparian strip regulators earlier in development. More generally, although not always significantly, ERS-cells tend to be more abundant in the lower nutrient availability condition.

These results demonstrate that this superimposed state is environmentally regulated. Variation in abiotic factors in these experiments, such as nutrient availability and osmotic pressure, can alter the transcriptional states of the root by modulating the proportion of cells that acquire the ERS, potentially to adjust the physiology of the root to different environmental conditions.

### ERS-encoded natural variation in root environmental responses

*Arabidopsis* accessions, which evolved under a variety of natural environments, offer a unique opportunity to study the molecular basis of the response to different environmental conditions. Since each of the replicate experiments performed for this study contained a mix of cells from two *Arabidopsis* accessions (Fig. 1A), we took advantage of this to explore the differences in cellular states, at the single-cell level, between five *Arabidopsis* accessions (Col-0, Ws-2, Cvi-0, C24, and Ler), and particularly in the ERS. The ERS was detected in all accessions and was taken into account when assigning identities to the cells, in order to identify cell state-specific DEGs in pairwise comparisons between accessions. The detected DEGs (average log fold-change ≥ 0.25, adjusted p-value < 0.05, t-test) are predominantly specific to one accession rather than shared between multiple (fig. S15), therefore we focused on the genes that are differentially expressed in one accession compared to the other four, and investigated the affected cell identities and associated biological processes (Fig. 6A, Table S5).

**Fig. 6.**
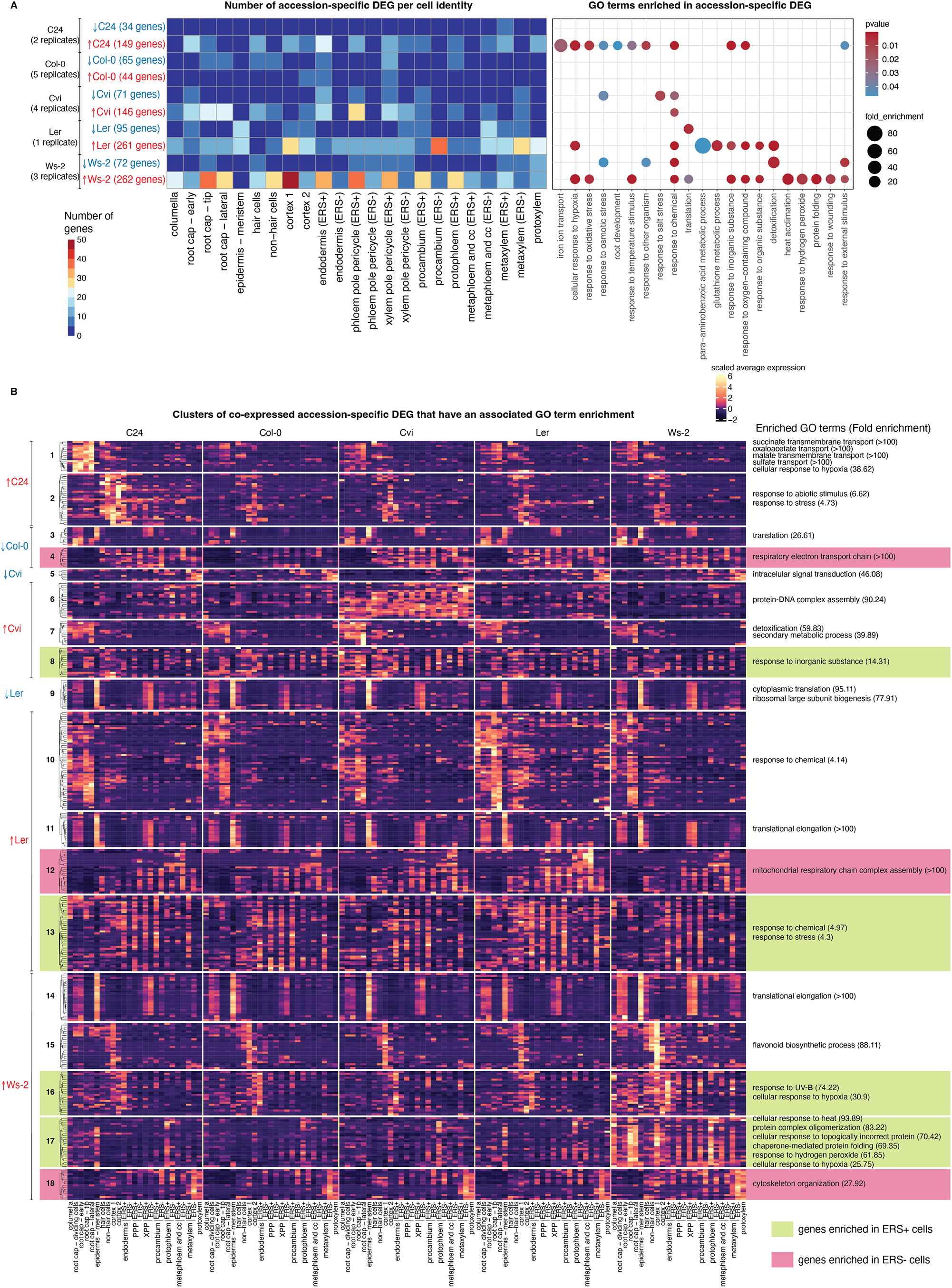
Natural variation in the environmentally responsive state. (**A**) Matrix showing the number of cell type-specific differentially expressed genes (DEGs) between Arabidopsis accessions and their associated GO terms. Matrix color indicates the number of DEGs, and arrow indicates the direction of differential expression in the indicated accession. (**B**) Heatmap showing the expression pattern of a subset in the clusters of co-expressed accession-specific DEGs (fig. S16) that have enriched GO terms.

Overall, the accession-specific DEGs are strongly associated with GO terms related to response to stresses and stimuli (Fig. 6A). The extent of differential expression differed between cell types; a larger proportion of genes was differentially expressed in the cells of the root cap and ERS+ cells in the stele, compared with other cell types. This was particularly apparent for upregulated DEGs in Ws-2, the category with the highest number of genes, which are associated with cellular processes linked to environmental responses such as heat acclimation. Furthermore, the number of DEGs is often higher in ERS+ cells as compared to ERS-cells of the same cell types, in multiple accession-specific comparisons. This further highlights that environmental responses are encoded in the ERS in the root.

We clustered the DEGs from each category based upon their expression pattern across the five accessions (fig. S16, A to J) and retained the eighteen clusters that had enriched GO terms (Fig. 6B). Among them, nine clusters correspond to genes that are enriched in a branch-specific manner, seven in ERS+ cells of multiple cell types and two in ERS-cells. While accession-specific DEGs enriched in ERS+ cells are almost always associated with responses to environmental stimuli, DEGs enriched in ERS-cells are involved in more structural and general cellular processes such as cytoskeleton organization and respiratory electron transport, again highlighting the strong association of the ERS to environmental responses.

*Arabidopsis* ecotypes have evolved in different environments, which is reflected in the genes that are differentially expressed among them. We show that a large proportion of accession-specific DEGs known to be involved in environmental responses are part of the superimposed ERS active state. This suggests that the new cell identity parameter we have defined is likely involved in the response to different natural environments.

## Discussion

Through deep characterization of root cell transcriptional states coupled to HCR spatial mapping, we have identified an alternate developmental trajectory for most root cell types, in particular the endodermis and the many stele cell types. This alternate path, which we term the environmentally responsive state (ERS), corresponds to a strong and specific transcriptional activity signature that is superimposed on a subset of the cells of multiple cell types simultaneously. Therefore, the identity of cells in the root tip can globally be characterized by three factors: 1) the lineage that defines the cell type, which is stable during development, 2) the developmental stage the cells are at, and 3) the presence or absence of an additional environmentally responsive state, which can vary based on environmental conditions

Although an ERS can be detected in multiple lineages, it is not equivalent for all cell types. Some of the genes are enriched in all ERS+ cells regardless of their developmental origin, but other ERS-specific genes are restricted to a few lineages, suggesting that there is a level of cell-type specificity in the acquisition of this transcriptional state, probably due to the underlying regulatory networks established in each cell type.

This cellular state (ERS+) is characterized by the expression of genes involved in the responses to various stimuli and stresses, the activation of the ABA signaling pathway - especially in the endodermis - as well as an enrichment for regulators and effectors of the circadian clock. While ABA signaling has been demonstrated to be important for the modulation of root growth in response to salt stress, particularly in the endodermis ^62,63^, and the circadian clock is known to be influenced by various types of external and internal inputs ^64^ as well as hormones such as ABA ^55^, here we provide the first demonstration that a network of pathways regulating the response to environmental cues are spatially restricted to a subset of gatekeeper cells of the root tip, and likely co-regulated. Future efforts will be required to identify the environmental cues and regulatory pathways that determine and control which cells will acquire the ERS.

We have demonstrated that the proportion of cells that acquire the ERS in each cell type can be modulated by environmental factors such as nutrient availability. Notably, we found a link between the branched states in the endodermis and a passage cell marker, suggesting that the regulation of the proportion of ERS+ cells could be a patterning event that pre-determines, early in development, the number of passage cells to be formed, thereby adapting the ability of the root to communicate with its environment. The discovery of this cellular state reveals new layers of complexity and cell specialization within the root, and opens the way to a better understanding of the role and integration of environmental stimuli in the modulation of root development.

## Acknowledgements

We thank Alex de Mendoza, Alexa Sadier, Olivier Hamant, Ian Small, and Justin Borevitz for critical feedback on this manuscript. We acknowledge the Centre for Microscopy, Characterisation & Analysis, The University of Western Australia, a facility funded by the University, State and Commonwealth Governments. We thank Chris Woodruff and Chengzhong Ye for assistance in developing genotype-based cell classification approaches. We thank Henry Löffler-Wirth for help with the use of the oposSOM package, and Harry Choi for sharing the HCR v3 protocol ahead of publication.

## Author Contributions

RL and MO conceived the study. MO and RL designed the experiments and analyses. MO performed the experiments and analyzed the data. TS performed the cell genotyping, data integration, and the analysis of bulk RNA-seq libraries. DT performed preliminary analyses of *Arabidopsis* accessions, and helped in establishing various analytical tools. SG contributed to the development of the cell genotyping method. JP and DP sequenced the libraries. JD, JJ, JW, and ML helped in conducting the pilot experiments. MO and RL wrote the manuscript. All authors approved of and contributed to the final version of the manuscript.

## Corresponding Authors

Correspondence and requests for materials should be addressed to Ryan Lister and Marina Oliva

## Data and Materials Availability

Raw and processed data has been deposited to GEO under accession number GSE198394.

## Supplementary Materials

Materials and Methods

Figures S1 to S16

Tables S1 to S5

References(*1-102*)

## Methods

### Plant material and growth conditions

Seeds from wild type *Arabidopsis thaliana* ecotypes Col-0, Ws-2, Cvi-0, c24 and Ler, and *scr-3* (Col-0 background, NASC stock code N3997) were sterilized and densely sown in row on 1x (or 0.5x, when testing the effect of nutrient concentration) Murashige and Skoog basal medium with vitamins (Austratec) plates supplemented with 1% sucrose (pH 5.7), and stratified in the dark at 4°C for two days. Plates were subsequently grown in long day conditions (16h light, 8h dark) at 22°C. 6-7 days after germination (dag), root tips (up to ∼3mm) were collected for protoplast isolation using sharp tweezers. Seedlings used in all experiments were grown in the exact same conditions, and collected and processed at the same time of the day, except for 2 out of the 10 replicates used to generate the single-cell atlas, which were performed in a different lab as part of a pilot experiment.

### Protoplast isolation and scRNA-seq

For each genotype, root tips were placed in 15ml of protoplasting solution ^65^ [1.25% w/v cellulase, 0.3% w/v macerozyme, 0.4M D-mannitol, 20mM MES (pH 5.7), 20mM KCl, 10mM CaCl2, 0.5% w/v BSA, and 5mM β-mercaptoethanol in ultrapure water] and incubated at room temperature for 1 hour on a tube rotator. The solution was subsequently filtered through a 70μm and a 40μm cell strainer (Falcon) to separate protoplasts from undissociated tissue. Cells were centrifuged for 5 minutes at 500 x g in a swinging bucket centrifuge. The pellet was resuspended in 15ml of protoplast resuspension buffer [0.4M D-mannitol, 20mM MES (pH 5.7), 20mM KCl, 10mM CaCl2, and 0.5% w/v BSA in ultrapure water] and centrifuged at 500 x g for 5 minutes. Cells were resuspended in the protoplast resuspension buffer, and filtered through a 35μm cell strainer Snap Cap (Falcon). The final concentration was adjusted to 900-1,200 cells/μl. Protoplasts from two *Arabdiopsis* accessions were mixed in equimolar quantities and barcoded with a Chromium Controller (10X Genomics). Libraries were generated with reagents from a Single Cell Gene Expression kit (10X Genomics, Single Cell 3’ Library kit) according to the manufacturer’s instructions. Libraries were sequenced either on an Illumina NextSeq 500 or a NovaSeq 6000, following recommendations of 10X Genomics.

### Preprocessing

Reads were mapped using a modified version of the 10x Genomics CellRanger pipeline (v1.3.0) ^66^. Internally, CellRanger uses STAR to perform read alignments, and the pipeline was modified to allow the specification of STAR insertion and deletion alignment penalties, as well as the minimum and maximum allowed intron size ^67^. Reads were mapped to a version of the *Arabidopsis* (TAIR10) genome with ambiguous base codes placed at SNP positions for the genotypes sequenced, as identified by the *Arabidopsis* 1001 genomes project ^68^. Reads were mapped with the following STAR parameters: --alignIntronMin=10, --alignIntronMax=5000, --scoreDelOpen=-1, --scoreDelBase=-1, --scoreInsOpen=-1, --scoreInsBase=-1.

### Cell genotyping

SNP UMIs were counted per cell using the countsnps command in the sctools package (https://github.com/timoast/sctools). Cells were then genotyped and cell doublets identified by performing density-based clustering on the SNP UMI counts for each genotype, implemented in the sctools genotype function. To classify cells at the border between those classified as “cell” and “background”, cells were further clustered in 30-dimensional principal component space using single cell gene expression data with an established graph-based clustering algorithm ^69–71^. Low UMI count droplets identified as “border” cells by sctools were given a “background” classification if they were assigned to a cluster with greater than 20% background droplets, otherwise they were classified as cells. This dimension reduction and clustering was performed using Seurat version 2.0.1 ^72,73^.

### Reference Atlas generation

#### Normalization and feature selection

The Seurat v3 R package was used for data normalization, cell filtering, batch correction, dimension reduction, and cell clustering ^16,72,73^. First, data were scaled to a total of 10,000 molecules and log-normalized. Cells with <200 expressed genes were removed, and genes expressed in <3 cells were removed. For each sample, the top 5,000 highly variable genes (HVGs) were detected using a variance-stabilizing transformation, using the FindVariableFeatures function in Seurat with the parameters selection.method=“vst” and nfeatures=5000. Genes affected by protoplast isolation were removed from this set of HVGs (see *“Identification of genes affected by protoplast isolation”* section below for more information).

#### Dataset integration

Technical differences between batches were removed using the data integration methods described by Stuart *et al*. ^*16*^. First, canonical correlation (CC) vectors were calculated between each pair of datasets, using the log-normalized expression of HVGs. Next, L2 normalization was applied to these CC vectors to correct for differences in scale between datasets. Anchor cells were then identified between each pair of datasets, using the first 30 CC vectors and the closest 5 neighbor cells. This was performed using the FindIntegrationAnchors function in Seurat v3.0.0 with the arguments anchor.features=3000, dims=1:30, eps=1. An integrated expression matrix was then produced using the IntegrateData function in Seurat v3.0.0 with default parameters. To produce an integrated expression matrix enabling the identification of common cell states across all experiments, we first ran this integration procedure using all anchor pairs identified between each pair of datasets. This matrix was used for downstream dimension reduction, clustering, and cell type classification. We also created a second integrated matrix that retained genotype-specific expression differences, while removing differences due to batch, but first filtering out cross-genotype anchor pairs from the integration procedure.

#### Cell clustering and visualization

The integrated expression matrix, described above, was used for all clustering and low-dimensional visualization of cells. The same set of 3,000 variable genes used for integration were used to calculate the top principal components, using the augmented implicitly restarted Lanczos bidiagonalization algorithm (IRLBA) ^74^. The top 50 principal components (PCs) were used to perform UMAP dimensionality reduction ^75^. Cell clustering was performed in the same 50-dimension PC space, using a graph based method as described previously ^69–71^. First, a k-nearest neighbor graph was constructed (k=30), and converted into a weighted shared nearest neighbor graph, with the graph weights representing the overlap between neighborhoods. Cells were then grouped into clusters through the identification of highly interconnected nodes in the graph, using the smart local moving algorithm ^76^. These steps were performed using the FindClusters function in Seurat v3.0.0.

### Identification of genes affected by protoplast isolation

Plants were prepared as for scRNA-seq, described above. Four replicates each of dissociated and of non-dissociated root tips were compared. For dissociated samples, root tips were placed in the protoplasting solution for 1 hour at RT, as for scRNA-seq. After this incubation, samples were centrifuged for 5 minutes at 500 x g, the supernatant was removed, and the tissues were snap-frozen in liquid nitrogen and ground into a powder. For non-dissociated samples, root tips were collected and directly snap-frozen and ground into powder in liquid nitrogen. Bulk RNA-seq libraries were generated using the TruSeq stranded mRNA kit (Illumina). Bulk RNA-seq libraries were sequenced on the Illumina HiSeq 1500 (single end, 101 cycles). RNA-seq read quality was assessed using FastQC and MultiQC ^77,78^. RNA-seq reads were trimmed with Trimmomatic v0.36 with the following parameters: ILLUMINACLIP:TruSeq3-SE.fa:2:30:10 LEADING:3 TRAILING:3 SLIDINGWINDOW:4:15 MINLEN:36 ^79^. Reads were aligned to the TAIR10 genome using STAR ^67^ with the following parameters: --alignIntronMax 5000, --alignIntronMin 10, --readFilesCommand zcat,--quantMode GeneCounts, --outSAMtype BAM SortedByCoordinate. Differentially expressed genes due to the cell dissociation were identified using the DESeq2 R package ^80^, with a Benjamini Hochberg adjusted p-value <0.001 and a log2 fold change in expression >2. This experiment was repeated twice, and genes that were identified as differentially expressed in both experiments were taken as the set of genes affected by protoplast isolation.

### Identification of markers from sorted cell populations

Raw sequencing data from Li et al. was downloaded from SRA (PRJNA323955) ^6^. Reads were trimmed to remove sequencing adapters and low quality bases using trimmomatic ^81^ with parameters ILLUMINACLIP:TruSeq2-PE.fa:2:30:10, LEADING:3, TRAILING:3, SLIDINGWINDOW:4:15, MINLEN:36. Genes were quantified using kallisto quant ^82^ with -b 100 to perform 100 bootstrap samples, using the TAIR10 CDS sequences. Differentially expressed genes specific for each FACS-sorted cell population were then identified using DESeq2 ^83^ with default parameters. Differentially expressed genes were then filtered to retain genes that were unique to one cell type, had a false discovery rate (Benjamin-Hochberg) <0.2, and a log2 fold change >1 compared to the mean expression across all other cell types.

### Whole-mount *in situ* HCR

#### Probe design

Candidate cluster markers for our reference atlas were generated using the FindAllMarkers function in Seurat, with the only.pos = TRUE parameter. To select the best target genes, cluster markers were filtered based on the following criteria: the gene should be detected almost only in the cluster(s) it is enriched in, and have a large portion of its coding sequence that doesn’t share homology with other Arabidopsis genes, to allow for design of highly specific probes for these sequences. Split probes for our target genes were designed and synthesized by Molecular Instruments (www.molecularinstruments.com).

#### Sample fixation and dehydration

The procedure is similar to the one described in ^84^. Roots were collected at 6 dag, as for scRNA-seq, and immediately placed in FAA [20% formaldehyde, 5% acetic acid, 50% ethanol, in ultrapure water] for 1 hour at RT. Samples were then dehydrated in successive 10 min washes at RT: once in 70% ethanol, once in 90% ethanol, twice in 100% ethanol. Samples were washed twice in 100% methanol and left in the second wash at -20°C overnight (or up to a few weeks).

#### Permeabilization, protease digestion

Roots were rehydrated in a series of 5 min washes at RT in 75%, 50%, 25% methanol in DPBS-T [0.1% Tween 20 in DPBS], and incubated in 1x cell wall digestion solution [10x stock: 250mg macerozyme, 250mg cellulase, 125mg pectolyase, 375mg pectinase in 50ml ultrapure water] in DPBS-T. After a wash in DPBS-T, samples were fixed in 10% (v/v) formaldehyde [37% solution, Sigma-Aldrich] in DPBS-T for 15 minutes at RT and washed twice in DPBS-T. Roots were incubated in 1ml Proteinase K buffer [0.1M Tris-HCl (pH 8), 50mM EDTA (pH 8) in ultrapure water] with 8μl Proteinase K (10mg/ml) for 15 min at 37°C. After two washes in DPBS-T, samples were fixed in 10% (v/v) formaldehyde [37% solution, Sigma-Aldrich] in DPBS-T for 15 minutes at RT and washed twice in DPBS-T.

#### Probe hybridization and amplification

We followed the third-generation *in situ* hybridization chain reaction procedure developed by Molecular Instruments, and described in ^15^, but the dextran sulfate used in the Amplification Buffer had to be replaced by a low molecular weight dextran sulfate in order to preserve the root structure.

Roots were pre-hybridized in the 30% probe hybridization buffer for 30 min at 37°C, then placed in the probe solution [1μl of probe mixture in 500μl of 30% probe hybridization buffer], and incubated at 37°C overnight.

Samples were washed twice in the 30% probe wash buffer for 30 min at 37°C, then washed twice in 5x SSCT for 10 min at RT. Samples were placed in the Amplification Buffer for 30min at RT. In the meantime, fluorescently labeled hairpins were prepared by snap cooling 10μl of 3μM stock in the hairpin storage buffer. We only used HCR amplifiers coupled to Alexa 647 fluorophores in this study. The hairpin solution was prepared by adding the snap-cooled hairpins to 500μl of amplification buffer. Samples were placed in the hairpin solution and incubated overnight at RT in the dark. They were subsequently washed three times in 5x SSCT for 20 min at RT.

#### Clearing

Samples were incubated in ClearSee ^17^ in the dark for three days.

#### Cell wall staining and sample mounting

Cell walls were stained by incubating samples in 1ml ClearSee + 2μl calcofluor solution [50mg/ml in DMSO] for 2 hours, in the dark. Samples were washed in ClearSee for 1 hour, in the dark, and mounted in ClearSee in a chamber made with a square coverslip and double-sided tape. The chamber was sealed using top coat nail polish, and the samples were imaged on the same day.

#### Imaging

Mounted roots were imaged using a Nikon Ti-E inverted motorized microscope with Nikon A1Si spectral detector confocal system, and a water immersion objective. Calcofluor white and the Alexa 647-coupled amplifiers were excited at 405 nm and 640 nm, respectively. Seven to eight images were taken per root along the differentiation axis. Images were processed using Fiji ^85^. Images of the same roots were stitched using the Grid/Collection stitching plugin ^86^, and the brightness/contrast was adjusted for each image based on the signal detected. 3D views of the root were obtained with the 3D viewer plugin ^87^, and optical sections with the Orthogonal Views method in Fiji.

### Identification of clusters 16 and 28

The first, *AT3G23830* (*RBGA4*), is detected in the meristem in various cell types including the epidermis (Fig. 1D). The cluster it is enriched in (cluster 16) is located between the differentiated epidermal clusters (hair cells and non-hair cells) and the early root cap clusters (Fig. 1B), suggesting that it represents meristematic cells of the epidermis. We have annotated this cluster as “epidermis - meristem”. The second, *AT3G25980* (*MAD2*), involved in cell division, is sporadically detected in different cell types in the meristematic region (Fig. 1D). Its associated cluster (cluster 28) in the dataset (Fig. 1B) appears to link the “epidermis - meristem” cluster to the “root cap - early” cluster. To test whether this cluster represents a specific population of mitotic cells in the early events of root cap formation, or an artifactual grouping of all dividing cells of the dataset, we regressed cell cycle effects out and looked at the influence on clustering (fig. S3, A to C). Cells that belong to cluster 28 still cluster together after regression and are also associated with early root cap and epidermis clusters, demonstrating that it corresponds to the known early formative cell division occurring in the epidermis/root cap lineage ^21^. We named this cluster “root cap - dividing cells”.

### Cell cycle regression

We used the CellCycleScoring function in Seurat to assign a cell cycle score to each cell of our reference atlas based on the expression of S phase marker genes [*E2Fc* (*AT1G47870*), *E2Fb* (*AT5G22220*), *DPa* (*AT5G02470*), *CDKG;2* (*AT1G67580*), *E2Fa* (*AT2G36010*), *DPb* (*AT5G03415*), *CYCA3;1* (*AT5G43080*), *CYCA3;2* (*AT1G47210*), *KRP5* (*AT3G24810*), *KRP2* (*AT3G50630*), *KRP6* (*AT3G19150*), *KRP3* (*AT5G48820*), *WEE* (*AT1G02970*), *KRP4* (*AT2G32710*), *CYCD4;2* (*AT5G10440*)], and of G2/M phase marker genes [*CYCB1;1* (*AT4G37490*), *CYCB2;1* (*AT2G17620*), *CDKB2;2* (*AT1G20930*), *CYCB3;1* (*AT1G16330*), *CYCA2;2* (*AT5G11300*), *CYCA2;4* (*AT1G80370*), *CYCA1;1* (*AT1G44110*), *CYCA2;1* (*AT5G25380*), *CDKD;1* (*AT1G73690*), *CDKB2;1* (*AT1G76540*), *CKS2* (*AT2G27970*), *CYCB1;2* (*AT5G06150*), *CYCD3;1* (*AT4G34160*), *CCS52B* (*AT5G13840*), *CYCA2;3* (*AT1G15570*), *CCS52A2* (*AT4G11920*), *CYCB2;3* (*AT1G20610*), *CYCB1;3* (*AT3G11520*)] obtained from ^88^. This score is used to provide a predicted classification of the cell cycle phase, and to regress out cell cycle effects using the vars.to.regress = c(“S.Score”, “G2M.Score”) in the function ScaleData.

After regression, the top 50 principal components were calculated and used for UMAP dimensionality reduction. Cell clustering was performed in the same 50-dimension PC space, using FindNeighbors and FindClusters functions in Seurat, as for the atlas.

### Pseudotime trajectory analyses

For the trajectory analyses, a new Seurat v2 object was created using Col-0 cells only and the batch-corrected data. Data was scaled and HVGs were detected using the FindVariableGenes function in Seurat v2 with the arguments x.low.cutoff = 0, and y.cutoff = -0.1. The genes affected by protoplast isolation we identified were removed from the set of variable genes. Principal components were computed and the top 30 PCs were used for UMAP dimensionality reduction, to generate the UMAP plot shown in Fig. 2 A and B.

The Col-0 dataset was then subsetted into eight objects corresponding to the developmental lineages: epidermis-root cap, cortex, endodermis, phloem pole pericycle, xylem pole pericycle, procambium, phloem, xylem. Each lineage dataset was then processed independently for the following steps.

#### “Stemness score” calculation

The diffusion pseudotime analysis required the identification of a cell that will be considered as t=0 at the starting point of the trajectory. Therefore, we developed an approach for identifying the “most meristematic” cell in each lineage.

We took advantage of the epidermis-root cap lineage having a cluster that is identified as meristematic, which links the epidermal clusters to the early root cap clusters. We first identified the markers specific to the meristematic state in the epidermis-root cap lineage, We generated a list of marker genes for all clusters of our reference atlas with the FindAllMarkers function with only.pos = TRUE argument. We defined the markers specific to the clusters 16, 9, and 12 as “early markers” for the meristematic region, and the markers specific to clusters 0, 1, 3, 4, 5, 8, 19, 25, 26, 27, and 28 as “late markers” for differentiated stages in the lineage. We expect genes common to the “early markers” and “late markers” to be specific to the lineage, but the genes specific to “early markers” to be specific to the meristematic zone. We therefore then subtracted the genes from the “early markers” that were also present in the “late markers” set, and used this meristematic gene list in the AddModuleScore (Seurat v2) to identify the cells that express this module of meristematic genes the highest in each lineage. We named this score the “stemness score” (plotted on UMAPs in fig. S4) and the cell with the highest score in each lineage is defined as t=0.

#### Sub-clustering and partition-based abstracted graph (PAGA)

HVGs were defined for each lineage object using the FindVariableGenes function (Seurat v2), with different x.low.cutoff and y.cutoff values. After removing genes affected by protoplast isolation from the HVGs, 40 PCs were computed and an elbow plot was generated using PCElbowPlot (Seurat v2) to determine the number of PCs that capture the majority of the variation in the data. This number of PCs varies between lineages (epidermis-root cap: 18; cortex: 8; endodermis: 8; XPP: 8; PPP: 7; phloem: 10; procambium: 10; xylem: 10).

Seurat lineage objects were then converted into AnnData objects using the Convert function (Seurat v2), in order to be processed with Scanpy ^20^. We first computed a neighborhood graph of observations, using the approximate nearest neighbor search within UMAP, with scanpy.pp.neighbors. The n_pcs argument was set, for each lineage, to the number of PCs determined above with the Elbow plot. Cells were subsequently clustered using the Louvain algorithm ^89–91^, with scanpy.tl.louvain. The resolution was adapted to each lineage, and, in some cases, some of the clusters had to be subsequently sub-clustered to increase PAGA clarity by using scanpy.tl.louvain with the restrict_to argument.

A partition-based abstracted graph (PAGA) was generated to quantify the connectivity between the lineage-specific clusters obtained above ^18^ using scanpy.tl.paga, and plotted with scanpy.pl.paga_compare, adjusting the threshold argument for each lineage. The PAGAs obtained for the eight lineages are shown in Fig. 2C.

#### Diffusion pseudotime

The cell of the lineage with the highest “stemness score” was annotated as the “root” cell. A diffusion map ^20,92,93^ was generated using scanpy.tl.diffmap, and the diffusion pseudotime ^18,19^ was calculated to infer the progression of cells through geodesic distance along the graph, using scanpy.tl.dpt. Both the sub-clustering and pseudotime assignment per cell were copied to the meta data of the original Col-0 Seurat object, and visualized in the UMAP plots in Fig. 2 A and B.

### Identification of genes with cell type- and pseudotime-dependent expression in the root cap (Fig. 2D)

A subsetted Seurat object (v3) containing only Col-0 columella and root cap cells was created. Genes that were detected in <175 cells were removed. Cells were categorized into bins of 100 cells, based on the developmental trajectory they belong to (columella, root cap - early, root cap - tip, and root cap - lateral) and their pseudotime assignment. The average expression of the detected genes was calculated for each bin using AverageExpression (Seurat v3). The obtained matrix of average gene expression per bin was scaled and subdivided into 12 clusters, using the kmeans function in R. The genes of one of the clusters, which did not show an interesting expression pattern, were removed from the scaled matrix. A heatmap showing the expression pattern of the remaining genes across all bins was generated using ComplexHeatmap in R ^94^.

### Assigning cell identities to published scRNA-seq datasets

Our reference atlas was subdivided into 82 clusters, and clusters were annotated based on the identities of the Col-0 cells obtained with the trajectory analyses (cell types and developmental branches).

Raw count matrices for Col-0 cells from six studies ^7–10,12,13^ were downloaded and used to create new Seurat objects. Cells with <200 expressed genes were removed, and genes expressed in <3 cells were removed. The data was log-normalized and the top 5,000 HVGs were detected using FindVariableFeatures (Seurat v3). Genes affected by protoplast isolation were removed from the HVGs.

Anchors between our reference atlas and the other datasets were identified using FindTransferAnchors in Seurat v3, specifying our atlas as the reference, using our genotype-corrected and batch-corrected integrated data as the reference assay, and the object we want to assign identities to as the query. The TransferData function (Seurat v3) was subsequently used to calculate, for each cell, a prediction score for each identity, based on the set of anchors previously identified and the annotations of the 82 clusters of the atlas. Cells were assigned the identity that has the highest prediction score.

### Identification of genes differentially expressed between branch 1 and branch 2

For our dataset and each of the published datasets, genes differentially between branch 1 and branch 2 of the endodermis, phloem pole pericycle, xylem pole pericycle, and procambium were identified using a t-test and the FindMarkers function in Seurat (v3), with the default average log fold-change threshold (0.25). A heatmap showing the average log fold-change (in natural log) for all the genes that were found differentially expressed in at least one of the comparisons was generated using ComplexHeatmap (fig. S5).

### UMI downsampling

Because the number of transcripts detected in branch 1 was systematically higher as compared to branch 2, we tested whether the branching events could be the result of coverage artefact by downsampling the UMIs to a similar rate in all cells. We chose to downsample the number of UMI to 4,000, which is the lowest median value of UMI per cell in the clusters of our dataset (cells that have a lower UMI count will not be downsampled). We used the SampleUMI function in Seurat (v4) to generate three independent downsampled count matrices, with the max.umi parameter set to 4,000. Each downsampled matrix was used to generate a new Seurat object that will be processed similarly to our original dataset (log-normalization, scaling, variable features, PCA, UMAP, clustering). An adjusted rand index was calculated using the adj.rand.index function of the fossil package in R, to compare cell clustering before and after UMI downsampling.

### Identification of gene co-expression modules using self-organizing maps (SOMs)

We used a SOM unsupervised machine learning approach to reduce the high dimensionality of the expression matrix and to cluster genes based on co-expression patterns. The matrix containing the scaled batch-corrected expression of genes across all Col-0 cells, except the genes affected by protoplast isolation, was used as an input for the oposSOM pipeline (v2.2) ^95,96^, via the opossom.new function with the following parameters: dim.1stLvlSom=60, feature.centralization=FALSE, sample.quantile.normalization=FALSE, activated.modules=list(“reporting”=FALSE, “primary.analysis”=TRUE, “sample.similarity.analysis” = FALSE, “geneset.analysis” =FALSE, “geneset.analysis.exact”=FALSE, “group.analysis”=FALSE, “difference.analysis”=FALSE. The SOM was then generated using the opossom.run function.

The metagenes in the SOM were subsequently clustered based on their Pearson correlation coefficients, using oposSOM. The following default parameters in the oposSOM source code for the correlation clustering were modified in order to optimize cluster specificity. In the pipeline.detectCorrelationModules function, the threshold for the correlation coefficient in the clustering was changed to 0.945 (default value: 0.9), and the minimal cluster length to 10 (default value: env$preferences$dim.1stLvlSom / 2). This resulted in smaller, and more specific co-expression modules. However, as a consequence, a number of small modules (<75 genes) were generated that had expression patterns almost identical to other bigger modules (>75 genes). We discarded these smaller modules and focused on the analysis of the 64 remaining modules, the composition of which is shown in Table S2. The average scaled expression per module, across all cells, was calculated using the scale.data matrix of the Col-0 Seurat object. Cells were categorized into bins of 100 cells based on their assigned developmental trajectory and pseudotime. An average of the average scaled module expression was calculated per bin, and plotted using ComplexHeatmap (Fig. 3A).

### UMAP plot of gene expression using schex

Because cells in UMAP plots are often plotted on top of each other, obscuring information and biasing interpretation, we used schex ^97^ in order to bin cells into hexagonal cells for visualizing gene or module expression. Hexagon bins were created for our Col-0 Seurat object, using the make_hexabin function with the parameter nbins = 150. Plot_hexbin_gene was subsequently used to visualize the scale expression of genes, or the calculated average scaled expression of genes from a co-expression module. The color scale was adjusted for each plot using scale_fill_viridis_c.

### Gene ontology enrichment per gene co-expression module

Gene Ontology (GO) enrichment analysis was performed using the PANTHER ^98^ Overrepresentation Test (Fisher’s exact test with Bonferroni correction). Genes were annotated using the GO Ontology database (biological process) ^99^, and all *Arabidopsis* genes were used as the reference gene list. Results with a corrected p-value <0.05 were considered significant.

PANTHER analysis results are sorted “hierarchically” in order to understand the hierarchical relations between enriched functional classes. Sorting is done only by the most specific subclass first, with its parent terms indented directly below it. These are all related classes in an ontology, and are often interpretable as a group rather than individually. Because the 64 modules had many enriched GO terms, some of which are related, only one term (the most specific one) per hierarchical group was selected per module. After this selection, a few very similar terms were filtered manually. The selected terms and the modules they are enriched in were plotted in Fig. 3B using ggplot2.

### Identification of candidate regulators using transcription factor (TF) target gene enrichment

Putative TF targets genes were inferred from the DAP-seq dataset ^53^. The ‘Motifs in peaks (P-value ≤ 1E-04)” BED file (http://neomorph.salk.edu/dap_web/pages/browse_table_aj.php) was used as a DNA binding database for the profiled TFs of the DAP-seq dataset.

500bp promoter coordinates for *Arabidopsis* genes were extracted using the promoters function in GenomicRanges ^100^, genes(TxDb.Athaliana.BioMart.plantsmart28) as an annotation for *Arabidopsis* genes, and the arguments upstream = 500, downstream = 50. For each TF, the binding coordinates were extracted from the Motif in Peaks .bed file and intersected with the promoter coordinates using the subsetByOverlaps function in GenomicRanges. Every gene that has a binding motif for the TF in the 500bp region upstream of its TSS is considered a putative target gene. For TFs that were not profiled in the DAP-seq studies but, according to the cisBP2 study ^54^, have a strong similarity in their DNA binding domains with another TF that was studied by DAP-seq, their target genes were considered to be the same as the TF profiled by DAP-seq.

In order to identify candidate cell-autonomous regulators in the gene co-expression modules we characterized, we looked for TFs in modules whose putative target genes are enriched in the same module the TF is expressed in. Only TFs whose putative targets could be inferred with the methodology described above could be tested. For each gene co-expression module, an enrichment test (hypergeometric test) was performed using the enrichment function of the bc3net package in R, with the genes in the module as the candidate genes, all *Arabidopsis* genes used as reference, and a list with the putative target genes for TFs in the module as the gene sets. A false discovery rate was used to correct for multiple testing and target genes were considered enriched in the module if the adjusted p-value was < 0.05. For the enrichment of ABRE at the vicinity of genes (Table S3) a similar test was performed, per module, using the combined putative targets of *AREB3* (*AT3G56850*) and *ABF2* (*AT1G45249*).

For candidate regulators of modules 47-49, the target genes list was reduced to the putative target genes that are part of the same co-expression module as the regulator. A network was generated using Cytoscape ^101^.

### Analysis of cell states in *scr-3* mutant root tips

#### UMAP generation and label transfer

A Seurat object was created with the batch-corrected *scr-3* data. The data was scaled and PCs computed with RunPCA. The top 30 PCs were used for UMAP dimensionality reduction. An identity was assigned to each cell using the same label transfer method in Seurat as the one we used for the published scRNA-seq datasets, described above. Anchors between our Col-0 object and the *scr-3* object were identified using FindTransferAnchors in Seurat v3, specifying our Col-0 object as the reference, and batch-corrected data as the reference assay. The TransferData function was subsequently used to predict an identity for each mutant cell, based on the set of anchors previously identified and using the developmental branched identified in the trajectory analysis to annotate the Col-0 reference cells.

#### Comparison of cell type composition between *scr-3* and wild type

Differences in cell type composition between *scr-3* and wild type Col-0 cells were assessed using the scProportionTest package in R ^102^. To test if the differences in cell proportion between two conditions isn’t simply a consequence of randomly sampling a number of cells, this package uses a monte-carlo/permutation test. Cells from both samples are pooled together and then randomly segregated back into two conditions, maintaining the original sample size. The proportional difference between the two conditions for each cell identity is calculated and compared to the observed proportional difference. This process is repeated 1,000 times, and the p-value represents the number of simulations where the simulated proportional difference was at least as extreme than observed (plus one) over the total number of simulations (plus one). This test was performed between the *scr-3* sample and each Col-0 replicate independently, which are approximately the same size. A cell type was considered significantly enriched or depleted if FDR < 0.05 and |log2(fold difference)|>0.58, in each comparison.

#### Early- and late-genes score in ground tissue lineage

The “early genes score” corresponds to the “stemness score” calculated with genes specific to the early stages of epidermis development in the epidermis-root cap lineage, but the AddModuleScore was applied to *scr-3* cells. The “late genes score” was similarly calculated using genes specific to late stages of epidermis differentiation in the epidermis-root cap lineage. This time, positive markers for clusters 3, 8, 19, 25, and 27 (late epidermal stages) of our reference atlas were identified and combined. From this list of markers we subtracted the genes that were specifically enriched in other clusters of the lineage (0, 1, 4, 5, 9, 12, 16, 26, 28). We used these “late genes” as features in the AddModuleScore function, applied to *scr-3* cells.

### Testing the effect of media composition on cell states

To test the effect of lowering nutrient concentration in the media, a set of plants grown on full (1x) MS or half (0.5x) MS were processed simultaneously in the exact same conditions, to avoid capturing other confounding effects. Two biological replicates were used per condition. Only Col-0 cells were used in these experiments, and therefore half of the doublets could not be removed based on SNP detection. However, doublets are expected to be present at the same rate in both conditions, hence not affecting the comparisons.

#### Data processing and cell identity assignment

A Seurat object with the four replicates was created. Cells with <200 expressed genes were removed, and genes expressed in <3 cells were removed. Data was log-normalized and scaled. The top 5,000 HVGs were detected using the FindVariableFeatures function in Seurat with the parameter nfeatures=5000. Genes affected by protoplast isolation were removed from this set of HVGs. PCs were computed and the top 30 PCs were used for UMAP dimensionality reduction.

Identities were assigned to cells using the FindTransferAnchors and TransferData as described above. Our Col-0 dataset with batch-corrected data, and the different cell types annotations with and without the ERS, were used as a reference.

#### Comparison of cell state composition

To compare the proportion of cells in each cell state between the two nutrient conditions we used the scProportionTest package in R as described above for *scr-3*. A test was performed for each pairwise comparison possible using the four replicates. Cell proportions were considered significantly different if FDR < 0.05.

#### Differential expression analysis

A differential expression test (t-test) was performed between each full MS replicate and each half MS replicate, per cell state (i.e. cell type with or without the ERS [ERS+, ERS-]), using the FindMarkers function in Seurat (v3), and the test.use=”t” parameter. The default log fold-change threshold used in this function is 0.25, and p-values are adjusted based on the Bonferroni correction.

To avoid taking into account variation in expression in cells where genes are very lowly expressed, we measured the average expression of all detected genes per cell state using the AverageExpression function, and also the mean of the average expression across all cell states. We then subsetted the DE results per cell state to keep only the genes whose average expression in the cell state is higher than the mean of the average expression across all cell states.

Genes that were systematically found up- or down-regulated in each of the four replicate comparisons were considered differentially expressed.

The average expression of DEGs per cell state and per condition was calculated, scaled, and plotted, together with results of the DE test, using ComplexHeatmap with the following parameters for hierarchical clustering of genes: clustering_methods_rows = “ward.D2”, row_split = 10.

#### GO term enrichment

A GO term enrichment was performed for each cluster of DEGs, using PANTHER and the GO Ontology database ^99^, as described above for the co-expression gene modules. Enriched terms were subsetted to keep only the most specific term per “hierarchical” group.

### Counting of hair cells

Seedlings were grown vertically on full (1x) or half (0.5x) MS plates, as for scRNA-seq experiments. Pictures of the roots were taken at 6 dag, using a stereomicroscope (n = 16 for full MS, n = 23 for half MS, from two different plates each). A portion corresponding to approximately 2.5mm from the tip was selected for each root picture, and root hairs were counted manually using the Cell Counter plugin in Fiji. A significant difference in root hair density between the two conditions was determined using a Wilcoxon Rank sum test (wilcox.test function) in R.

Note that the seedlings used for this quantification were in the “top” row of the plate (seeds are sown in two rows for scRNA-seq experiments). Roots of seedlings grown in the bottom row tend to have fewer root hairs (in both conditions).

### *Arabidopsis* accessions comparison

Our reference atlas was subdivided into 82 clusters, and clusters were annotated based on the cell state annotation of Col-0 cells obtained with the trajectory analyses (cell types with or without the ERS [ERS+, ERS-]).

#### Differential expression analysis

The dataset is composed of different numbers of replicates per accession. For each possible inter-accession replicate pair, a differential expression test (t-test) was performed, per cell state, using the FindMarkers function in Seurat (v3), the test.use=”t” parameter and default log fold-change and p-value cutoffs. To avoid taking into account variation in expression in cells where genes are very lowly expressed, we measured the average expression of all detected genes per cell state, in Col-0 cells, using the AverageExpression function, and also the mean of the average expression across all cell states. We then subsetted the DE results per cell state to keep only the genes whose average expression in the cell state is higher than the mean of average expression across all cell states. Genes were considered DE between two accessions only if they were systematically up- or down-regulated in all possible replicate pairwise comparisons.

For a gene to be called an accession-specific DEG, it had to be systematically DE in a given accession compared to the other four, but not be DE in any pairwise comparison between the remaining four accessions. Similarly, DEGs specific to two accessions had to be systematically DE in the two accessions as compared to the other three, but not be DE in any pairwise comparison between the remaining three accessions. As shown in fig. S15, most DEGs were specific to one accession, rather than two, hence we subsequently focused on those specific to one accession.

#### Gene clustering

For each category (up- or down-regulated in a specific accession), the average expression per DEGs per cell state and per accession was calculated, scaled, and plotted using ComplexHeatmap. The kmeans function was used to cluster DEGs based on their expression pattern, and the number of clusters was adjusted for each category.

#### GO term enrichment

A GO term enrichment was performed for each category of accession-specific DEGs (Fig. 6A) and for each cluster per category (fig. S16), using PANTHER and the GO Ontology database ^99^, as described above for the co-expression gene modules. Enriched terms were subsetted to keep only the most specific term per “hierarchical” group.

Only the clusters that have at least one enriched GO term were displayed in Fig. 6B.

