## Supplementary Figures for "An environmentally responsive transcriptional state modulates cell identities during root development"

Supplementary Figure 1

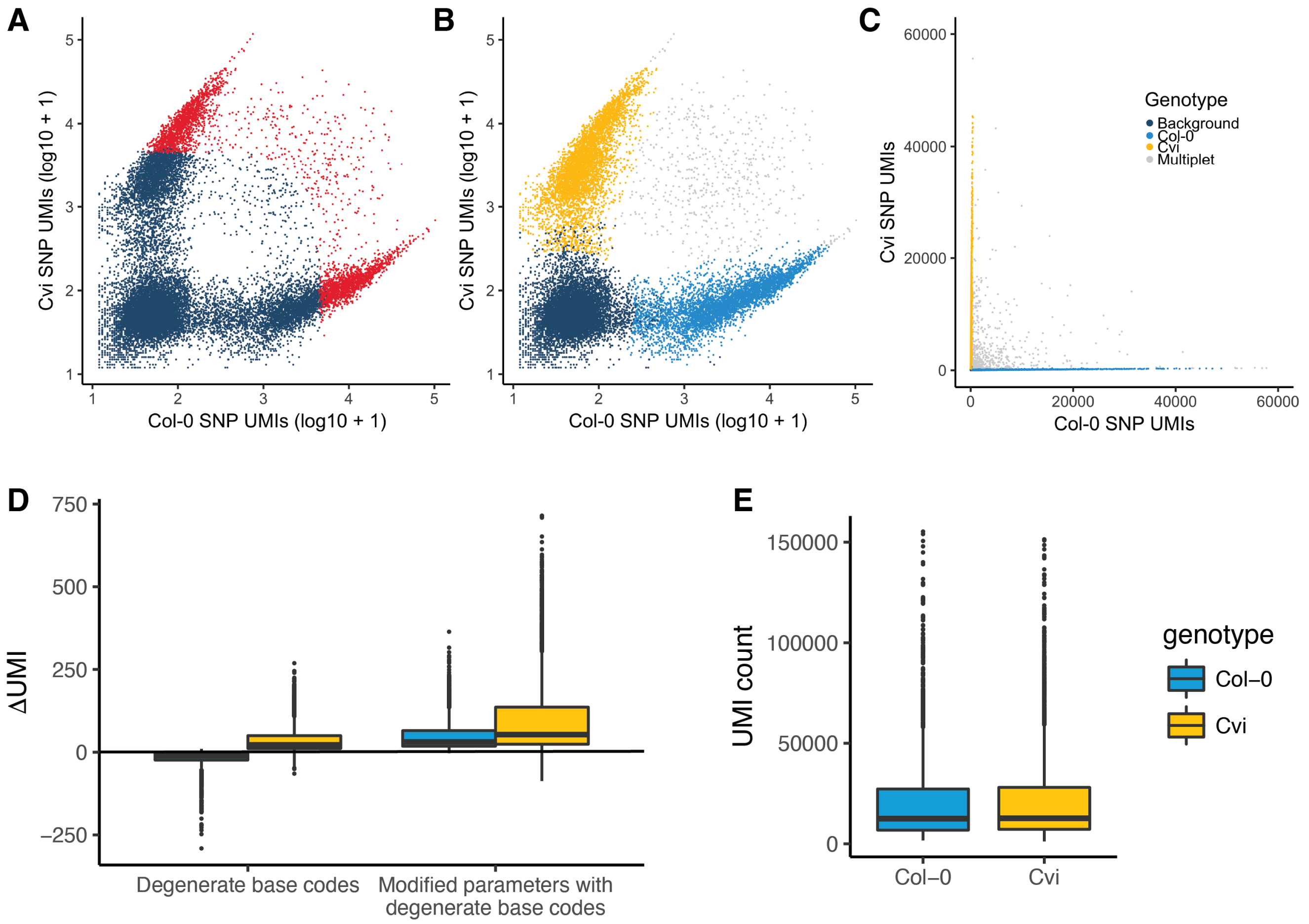

**Fig. S1. Genotyping of single cells based on single-cell SNP UMI counts**

(**A**) Single nucleotide polymorphism (SNP) unique molecular index (UMI) counts for Col-0 and Cvi specific variants for each scale (log10 scale). Color indicates the cell or non-cell classifications produced by the standard 10x Genomics CellRanger software, with empty droplets in blue and cells in red. (**B**) As for (A), but colored by genotype classification. (**C**) As for (B), using a linear scale. (**D**) Change in total UMI counts per cell when mapping to different genome versions, for Col-0 and Cvi cells. Introducing degenerate base codes at SNP positions results in a greater increase in total UMI counts for Cvi cells relative to Col-0 cells. (**E**) Distribution of total UMI counts per cell for Col-0 and Cvi cells.

Supplementary Figure 2

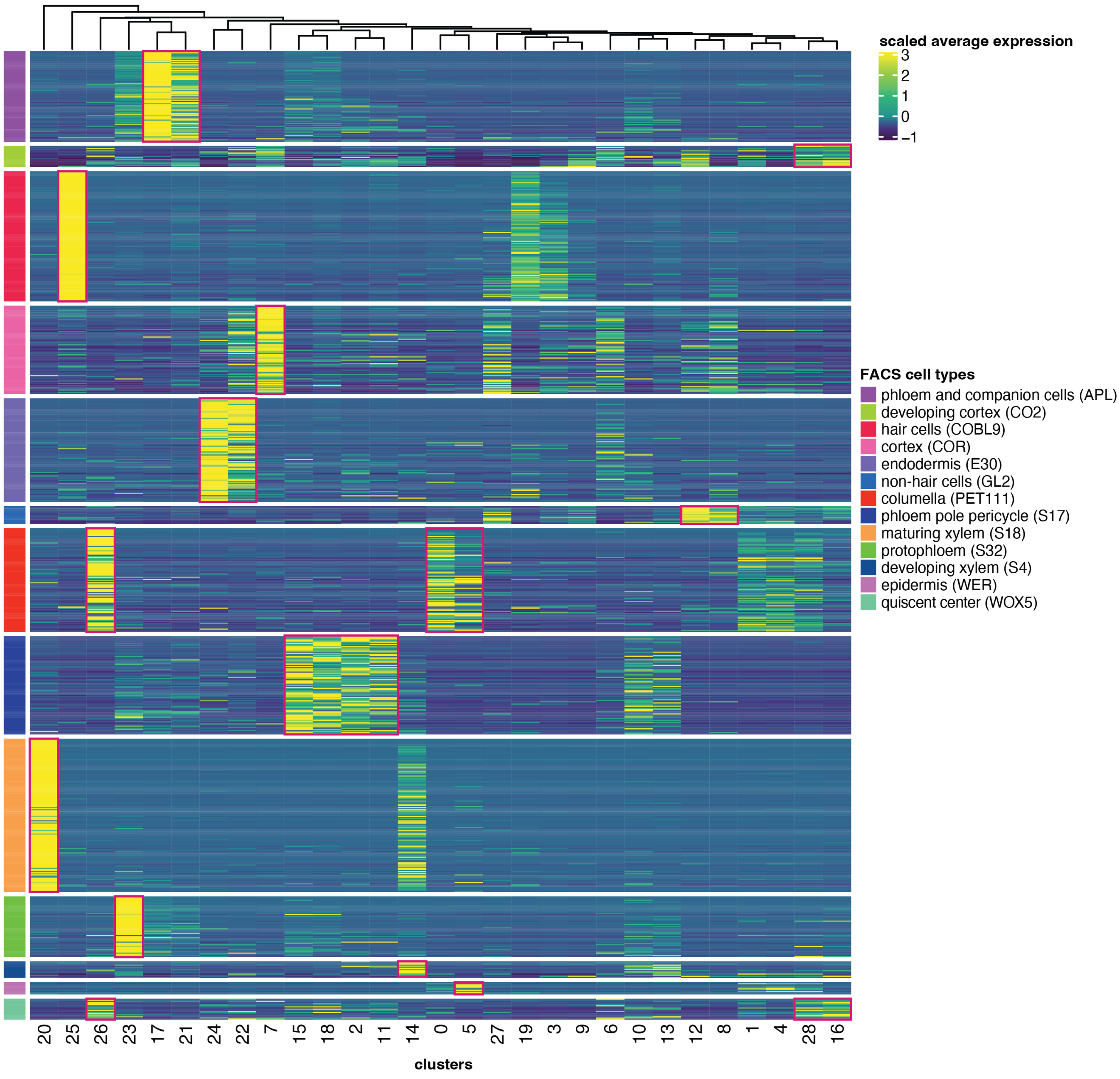

**Fig. S2. Expression of markers from cell type-specific bulk RNA-seq in the single cell dataset from this study**

Heatmap showing the expression of the cell type-specific markers identified from published RNA-seq of sorted cell populations (*6*) in the clusters of our reference atlas. Red rectangles indicate a cluster-specific enrichment of some of the marker genes. Not all cell types of the roots are represented with specific markers that were identified from the sorted populations, and multiple clusters do not have a matching set of markers identified in the sorted cell population data.

### Supplementary Figure 3

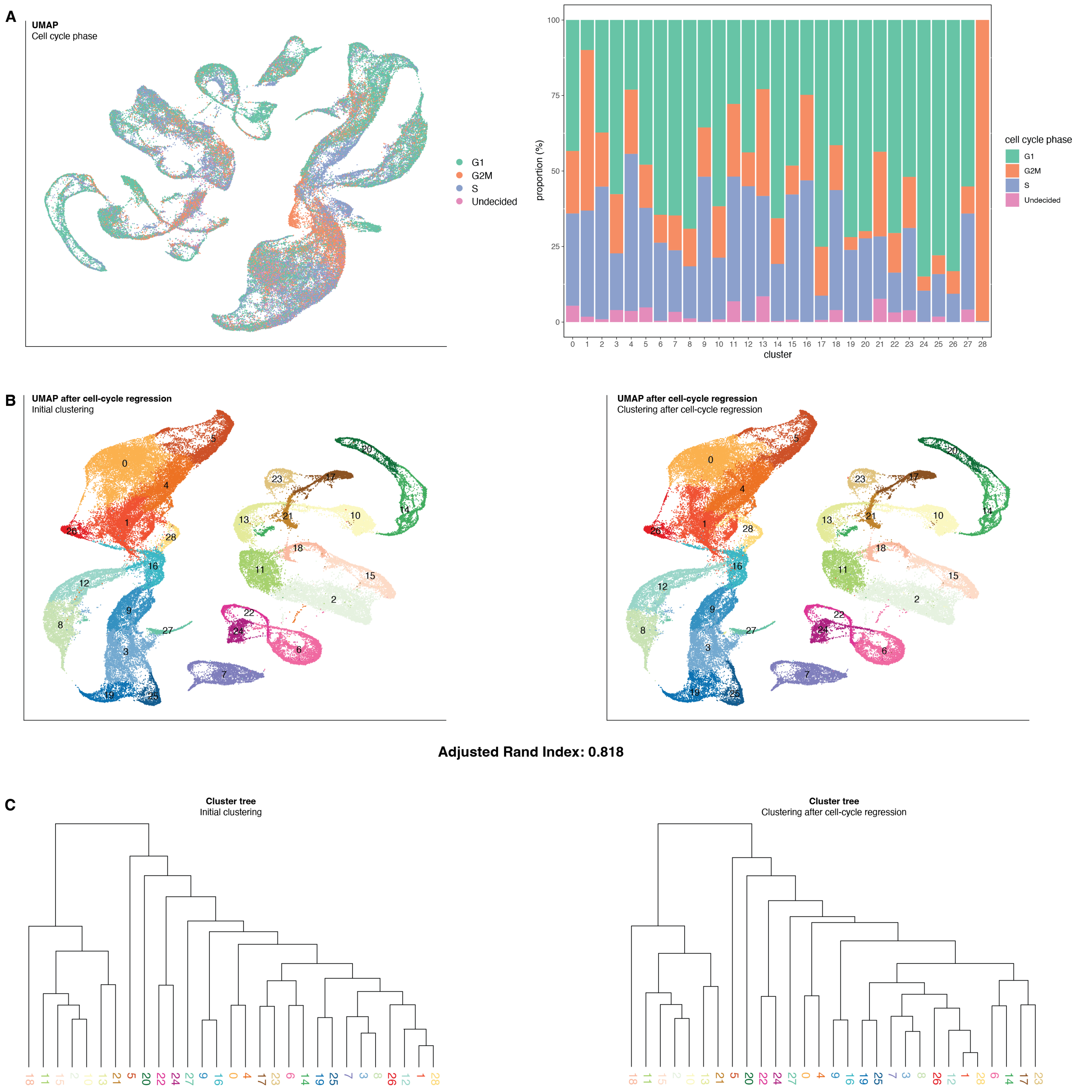

**Fig. S3. Cell clustering is not driven by the cell cycle**

(**A**) UMAP showing the predicted cell cycle stage of cells in the reference atlas from this study. The barplot shows the proportion of cells in each stage per cluster. (**B**) UMAP of the reference atlas from this study after regressing cell cycle effects. Colors show the clustering of cells before (left) and after (right) the regression. The Adjusted Rand Index of 0.818 comparing clusters before and after regression show that the effect of the cell cycle regression is minimal. (**C**) Phylogenetic tree depicting similarity of clusters before (left) and after (right) regression.

Supplementary Figure 4

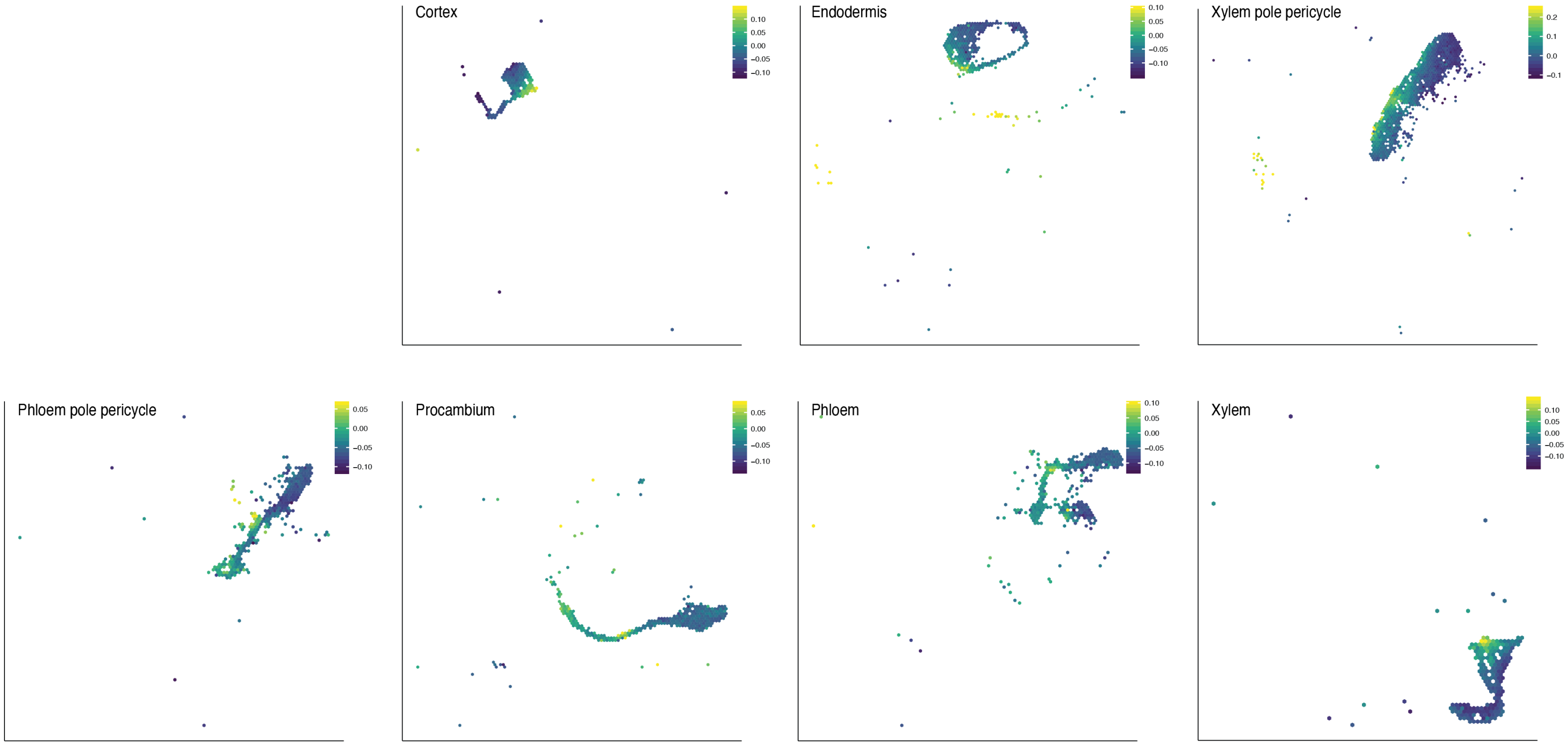

**Fig. S4. Identification of meristematic cells as starting points in each lineage trajectory**

UMAP plot of each Col-0 developmental lineage showing the “stemness score” per cell. This score is calculated from the expression of meristematic genes identified in the epidermis-root cap lineage (see Methods). The higher the score, the more likely the cells are close to the meristem. Cells were grouped in hexagonal bins using schex[97].

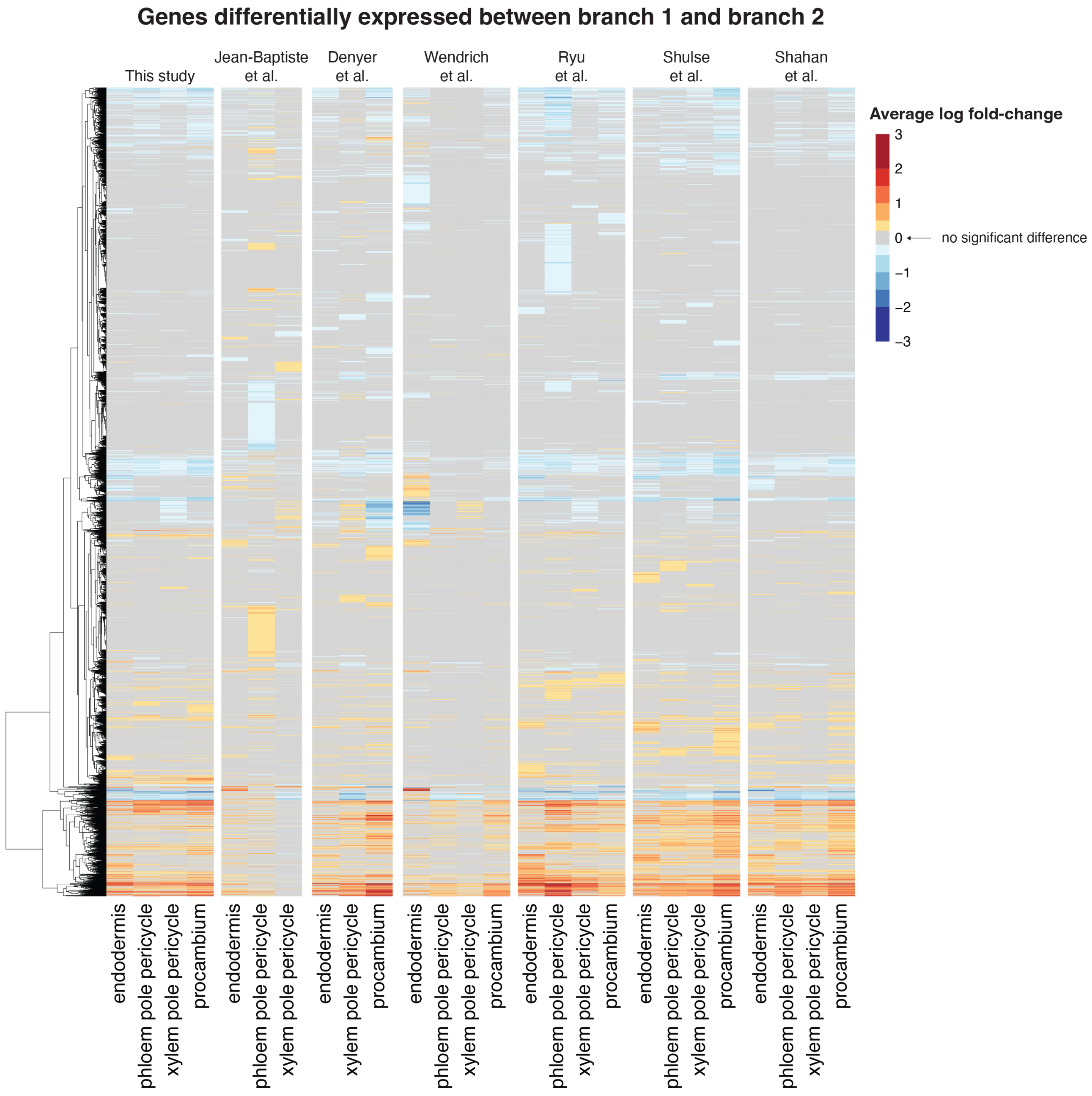
Genes differentially expressed between branch 1 and branch 2
This study
Jean-Baptiste et al.
Denyer et al.
Wendrich et al.
Ryu et al.
Shulse et al.
Shahan et al.
Average log fold-change
3
2
1
0
−1
−2
−3
no significant difference
endodermis
phloem pole pericycle
xylem pole pericycle
procambium

**Fig. S5. Cells of the alternative branches have similar signatures in all scRNA-seq datasets**

Heatmap showing the average log fold-change between branch 1 and branch 2, in the endodermis, phloem pole pericycle, xylem pole pericycle, and procambium. Our analyses identify all these genes as differentially expressed between branch 1 and branch 2, in at least one of the cell types, in at least one of the studies.

Supplementary Figure 6 (1/3)

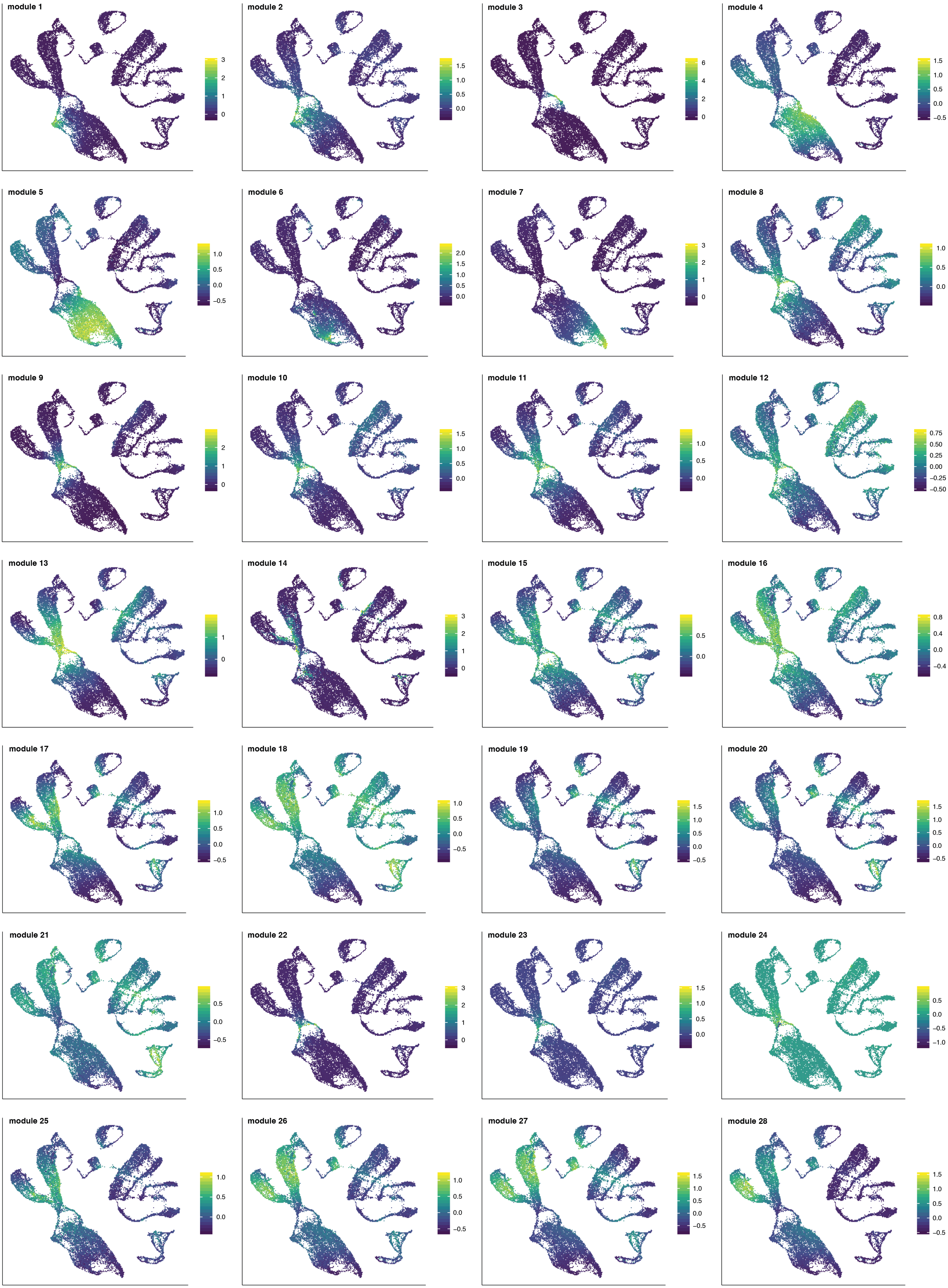

Supplementary Figure 6 (2/3)

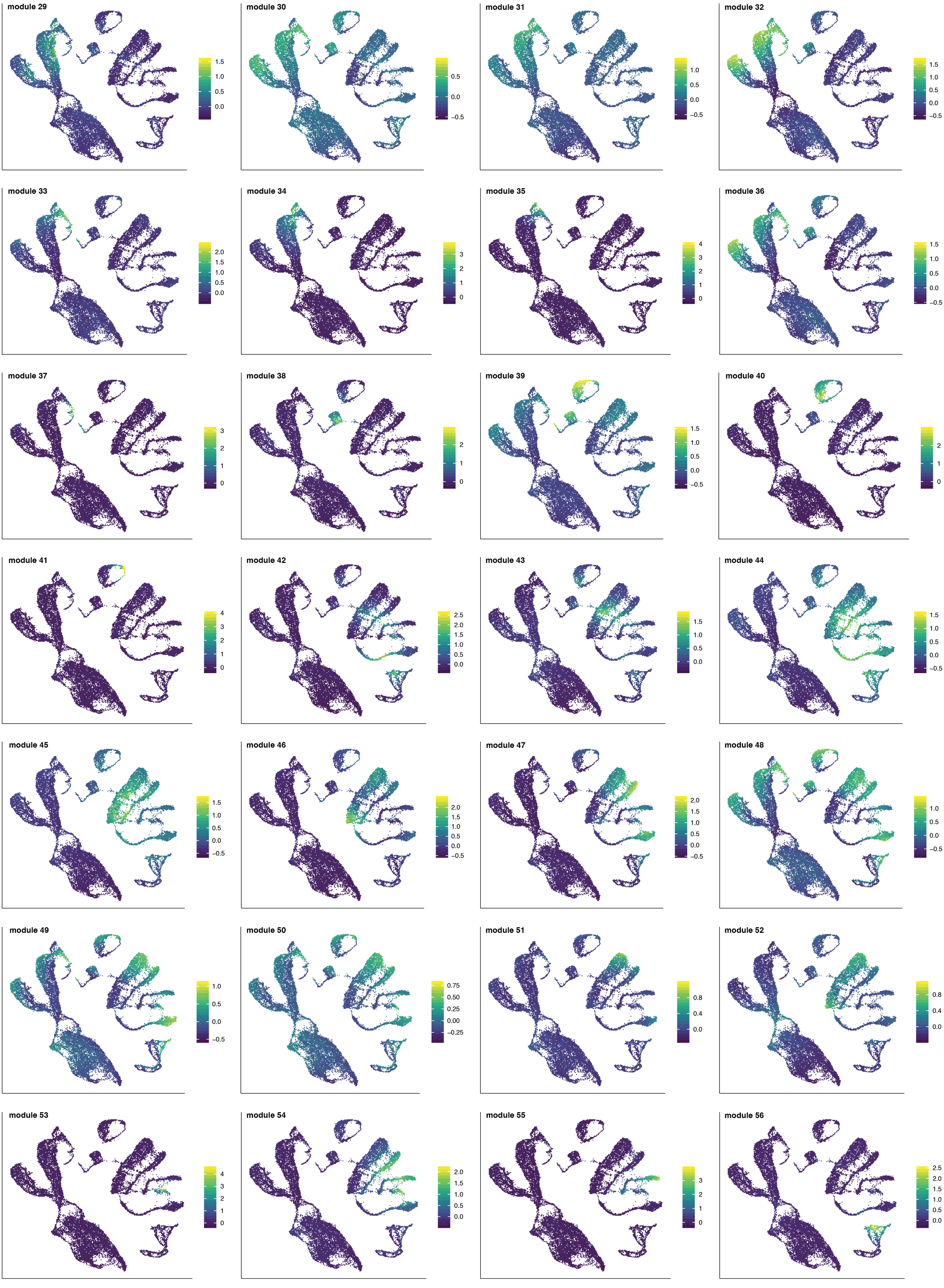

Supplementary Figure 6 (3/3)

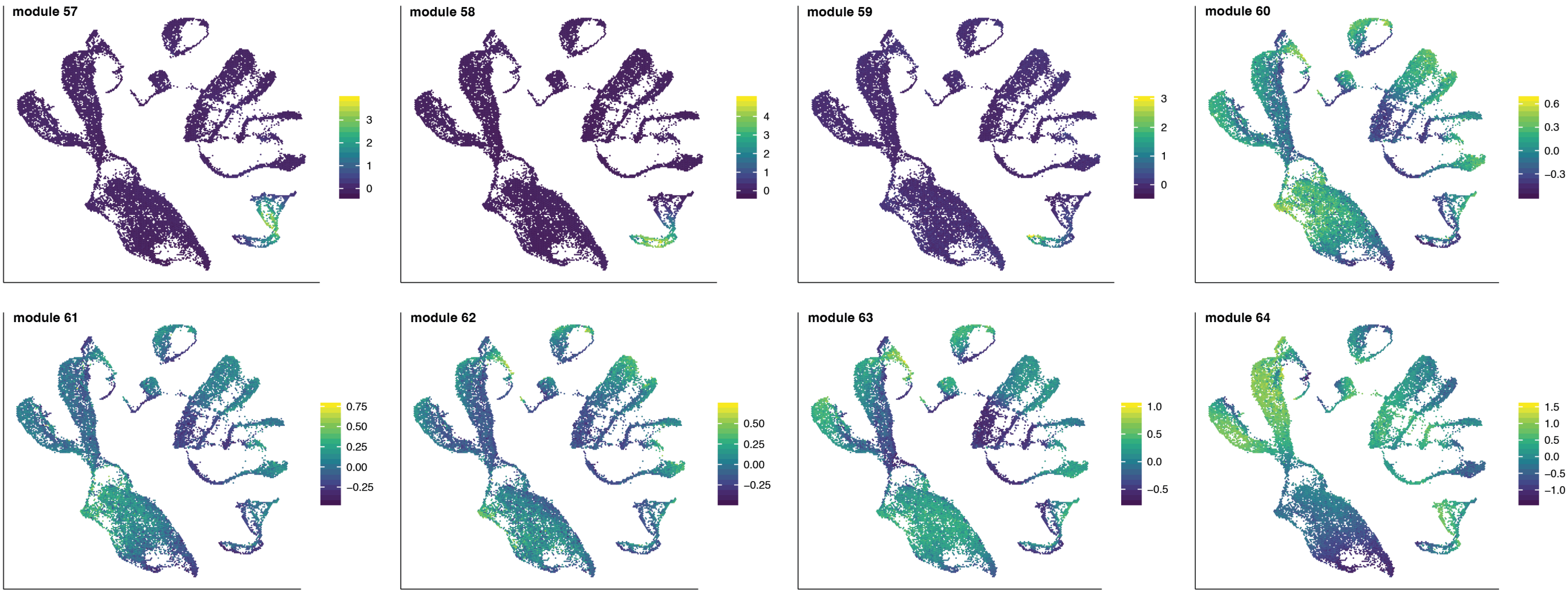

**Fig. S6. Expression patterns of the co-expression gene modules**
UMAPs showing the average scaled expression of genes per module (defined in Fig. 3). Cells are grouped in hexagonal bins using schex[97].

### Supplementary Figure 7

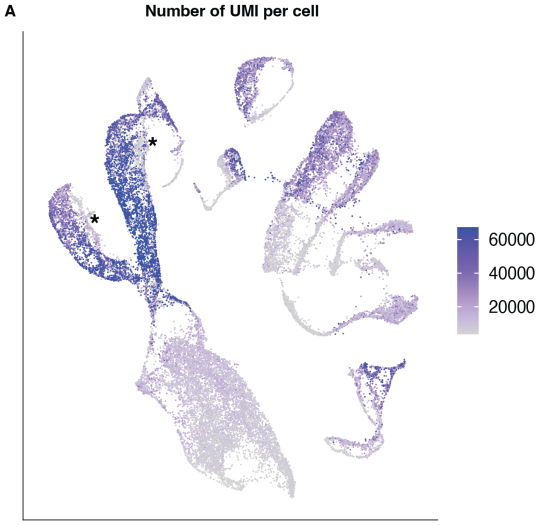


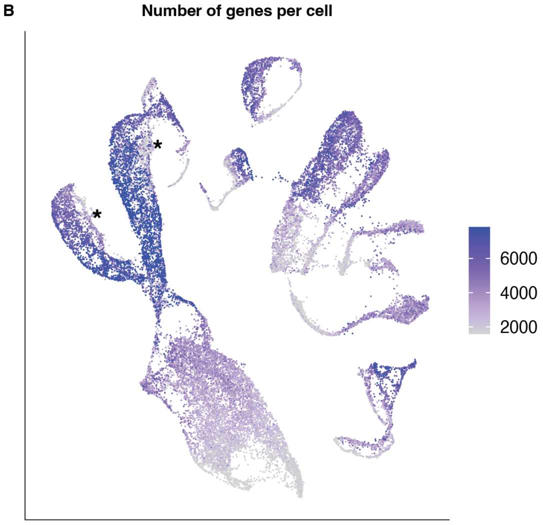


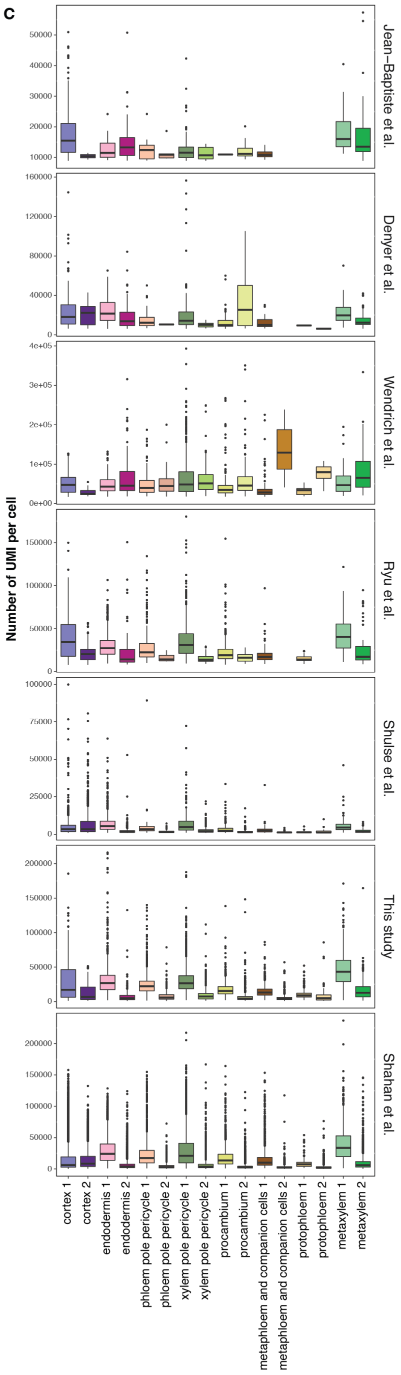

**Fig. S7. Branch 1 exhibits high transcript abundance as compared to branch 2, in multiple cell types**

(**A**) UMAP showing the distribution of the number of UMIs per Col-0 cell. (**B**) UMAP showing the distribution of the number of genes detected per Col-0 cell. The asterisks in (A) and (B) indicate a small population in the epidermal cell types that have a lower number of UMIs and genes. (**C**) Boxplots showing the distribution of the number of UMIs in cells assigned as one of the two alternate developmental branches, in multiple cell types, for each scRNA-seq study.

Supplementary Figure 8

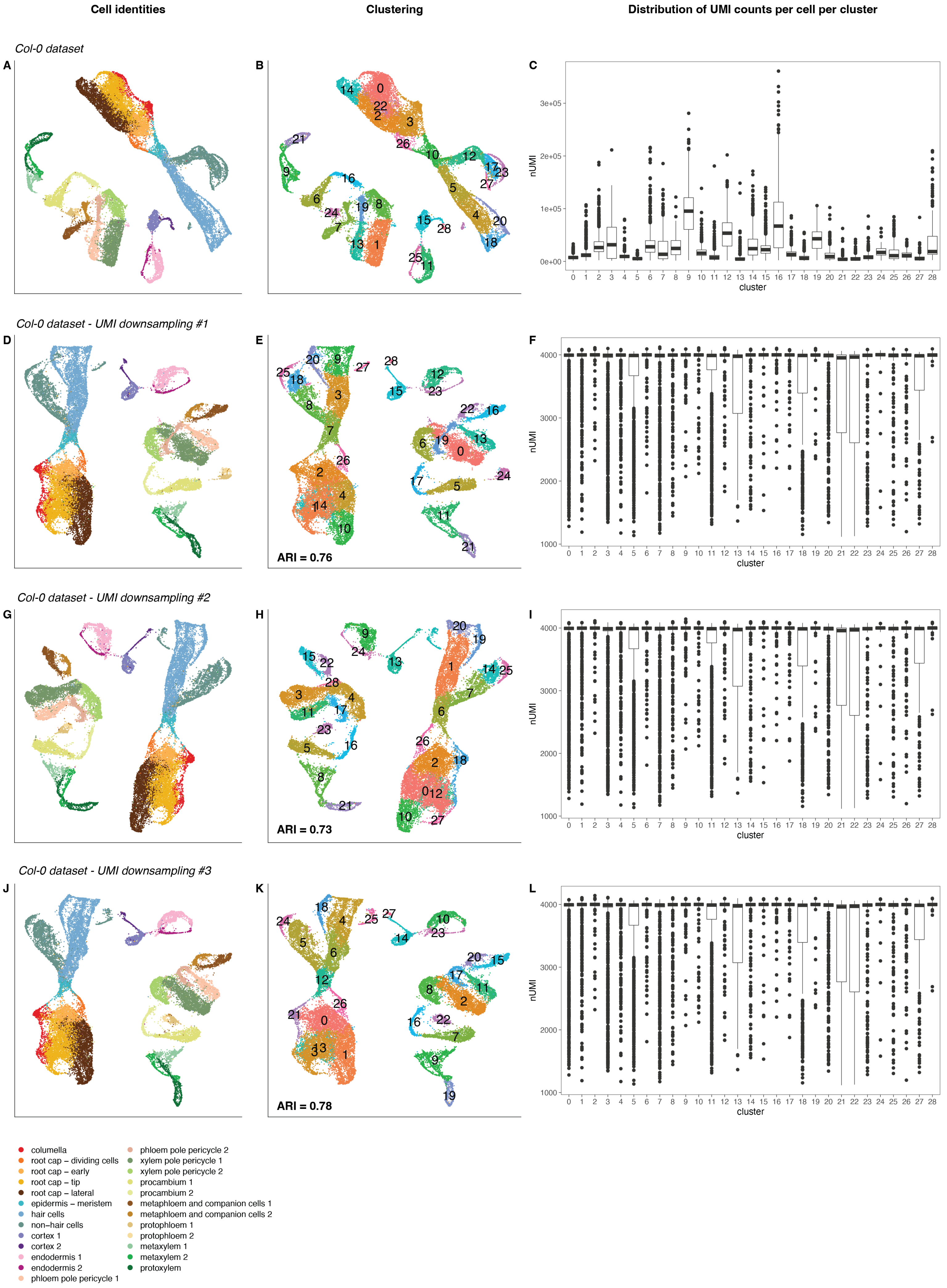

**Fig. S8 Branching events are not an artefact caused by coverage**

UMAPs showing cell identities (A, D, G, J) as determined in (Fig. 2), cell clustering (B, E, H, K), and the distribution of the number of UMIs per cell per cluster (C, F, I, L) in the Col-0 dataset before (A-C) and after 3 independent UMI downsampling computations (see Methods). The adjusted rand index (ARI) was calculated to compare clusters after downsampling (E,H,K) to the clusters before downsampling (B). After UMI downsampling, cells are still grouped by their branch identities (D,G,J) and clusters are very similar to the ones before downsampling (all ARI > 0.73), demonstrating that the branches are not an artefact caused by coverage (number of UMIs per cell).

Supplementary Figure 9

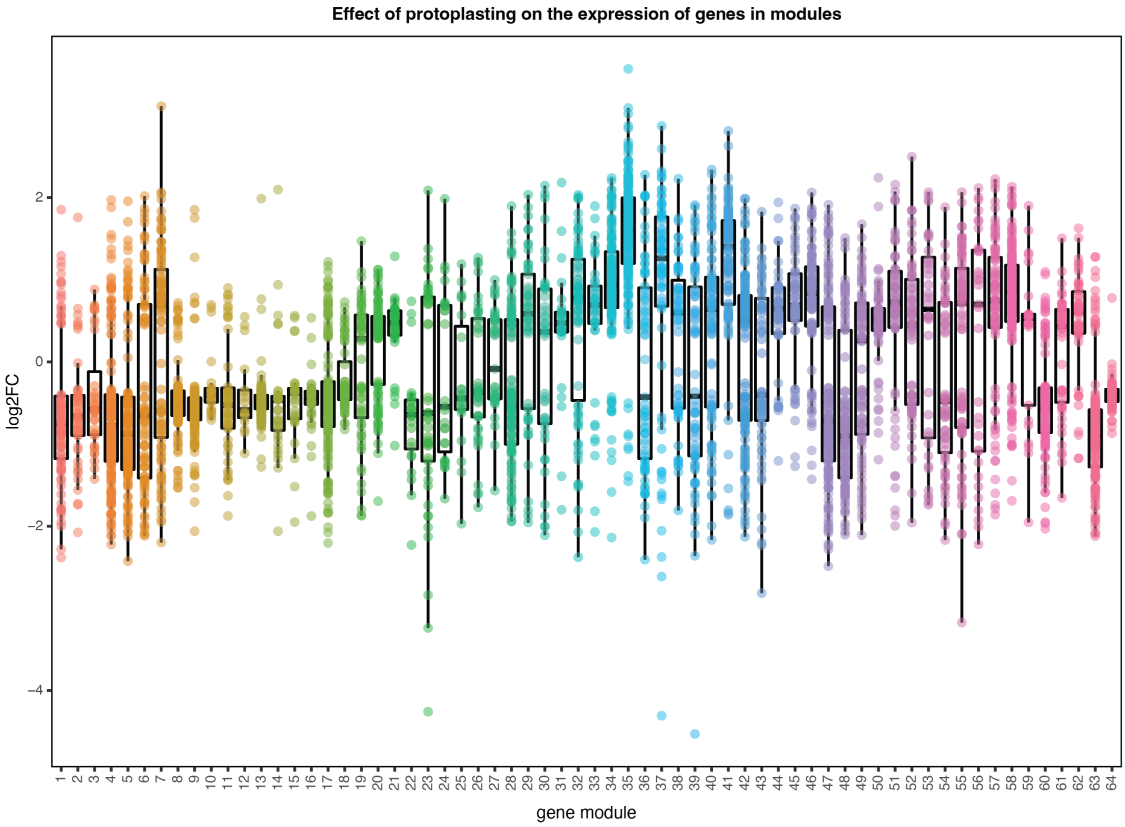

**Fig. S9. The branch-specific modules are not a result of cell dissociation**
Boxplots showing the distribution of the log2 fold-change in expression for the genes of the co-expression modules (Fig. 3A) as a result of protoplast isolation. Branch 1-specific modules 47-51 are not particularly up- or down-regulated in cells during protoplast isolation.

**Network of terms associated with the "response to hormone" genes of the branch-specific modules 47-51**

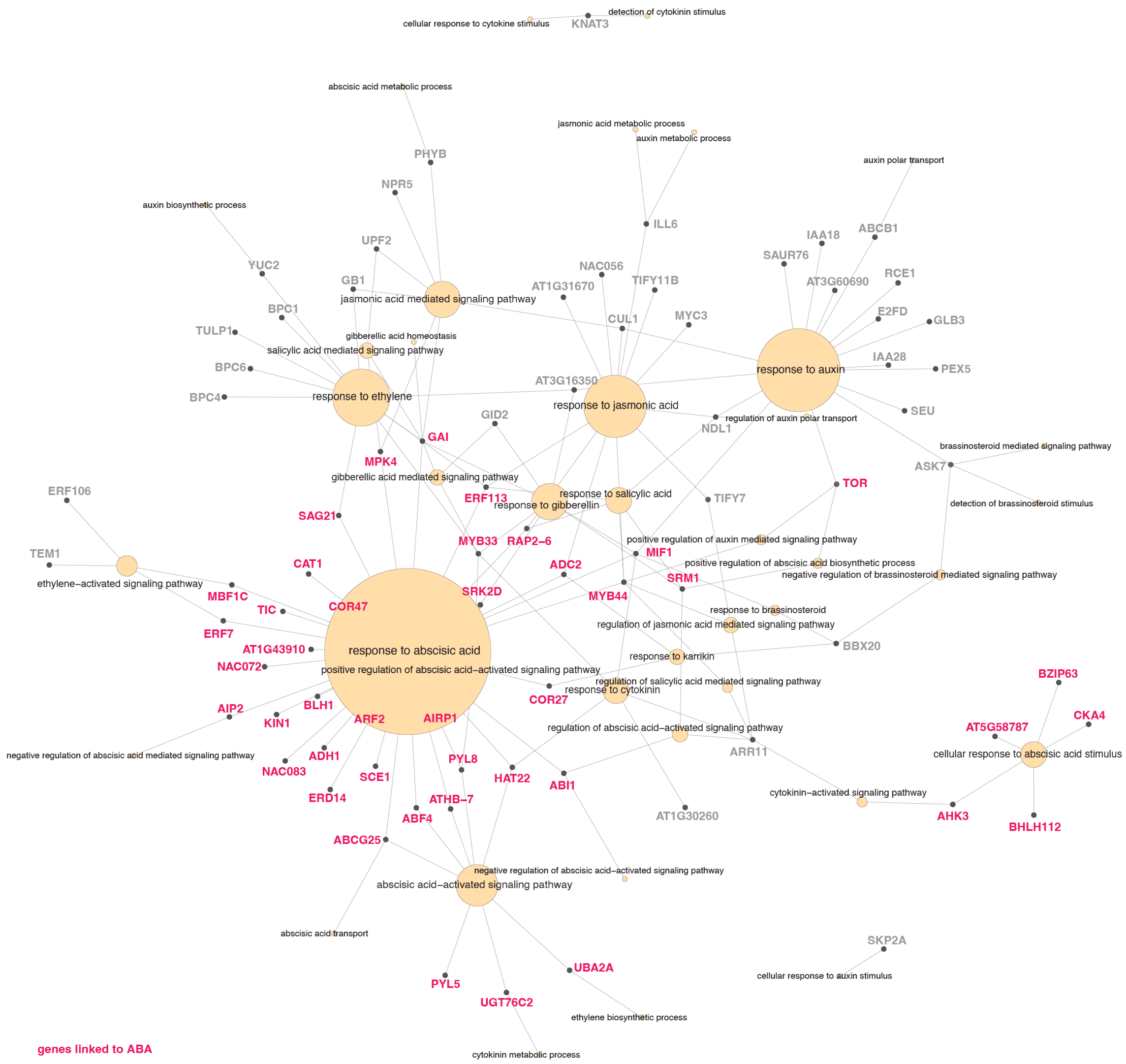

**Fig. S10. The branch 1 state is characterized by an ABA-responsive transcriptional signature**

Network showing all the genes (black dots) annotated as involved in *“response to hormone”* in the branch 1-specific modules 47-51, and all their associated GO terms linked to plant hormones (yellow circles). The size of the circles is proportional to the number of genes linked to them. Genes that are associated with terms involving ABA are colored in pink. More than 50% of the genes (42/78) are linked to ABA.

### Supplementary Figure 11

#### Circadian clock / Regulation of circadian clock

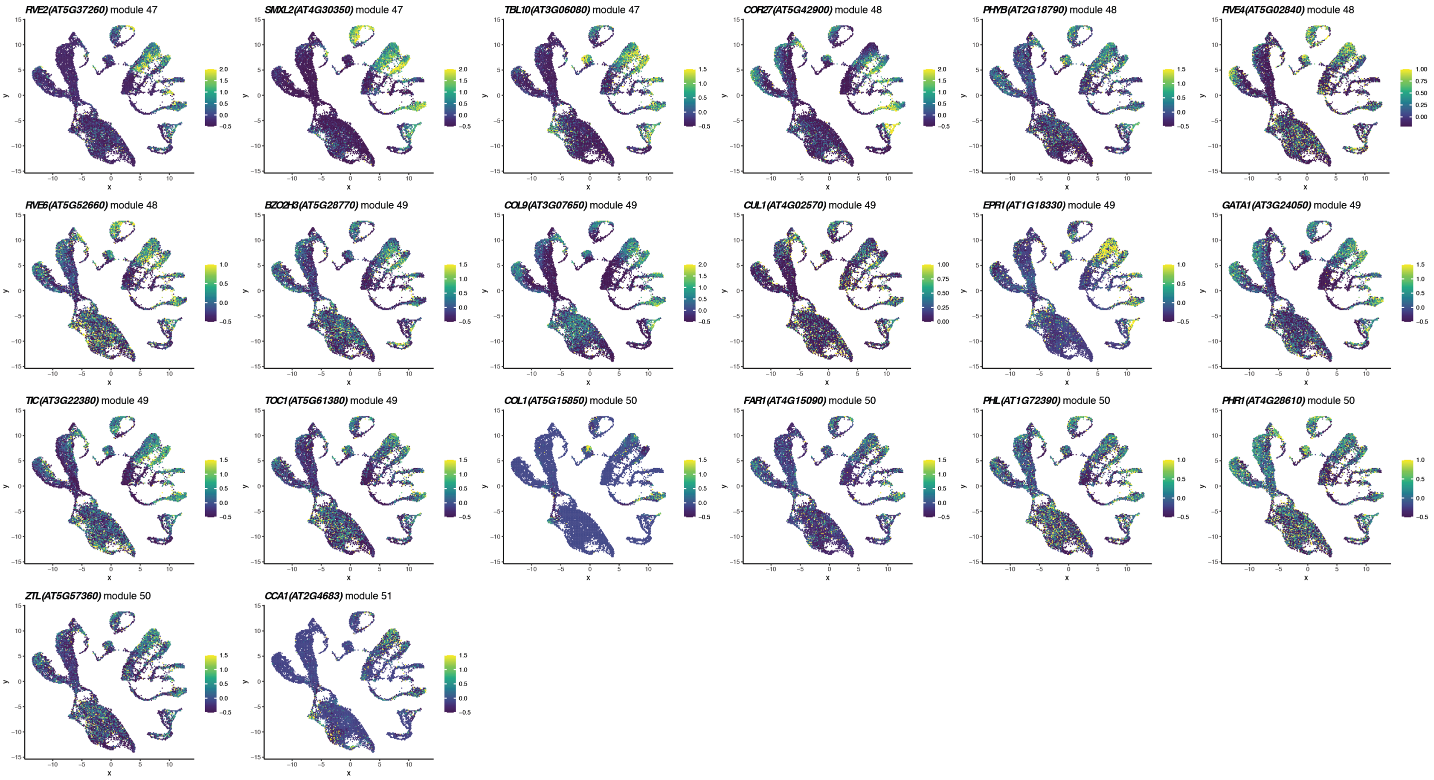


#### ABA-activated signaling pathway

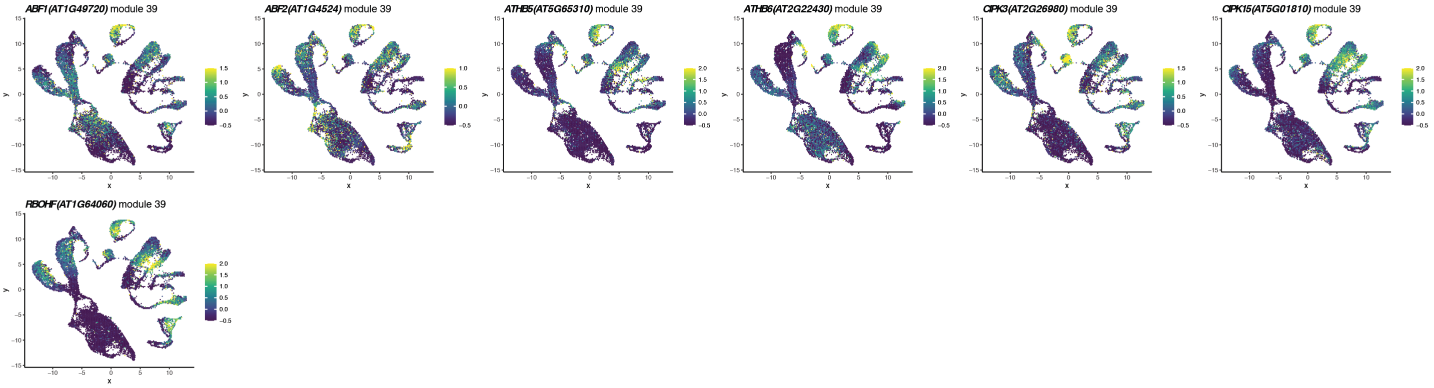


#### Regulation of ABA-activated signaling pathway

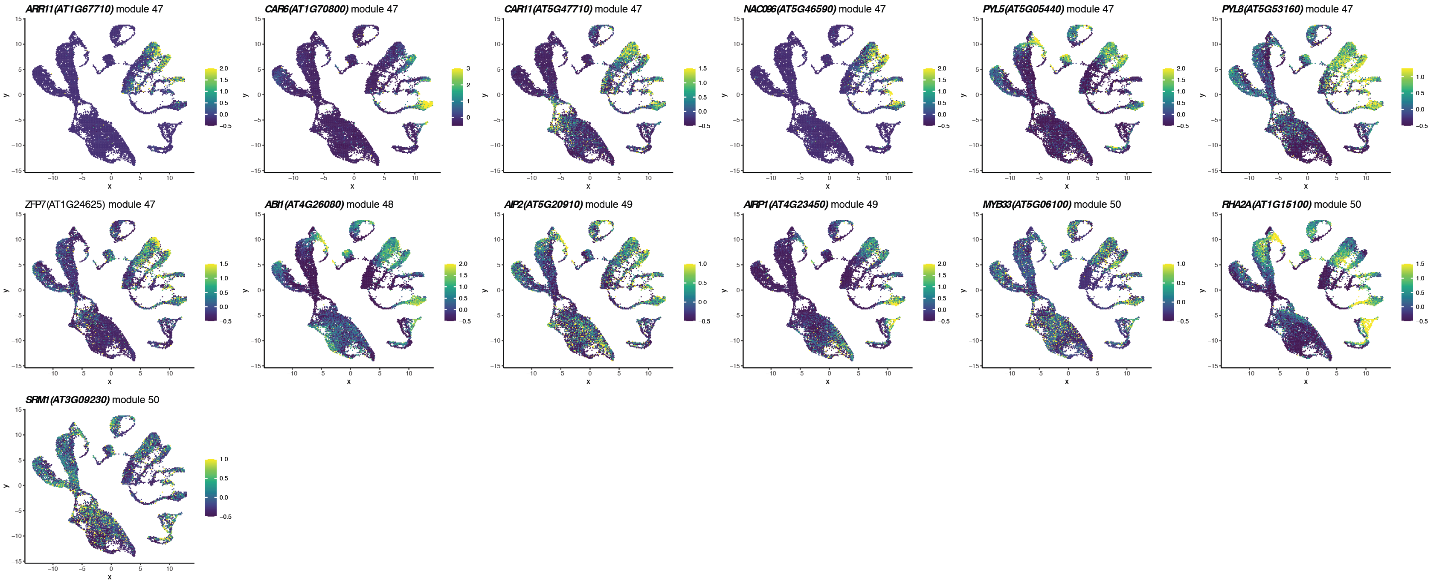

**Fig. S11. Expression pattern of the genes in branch 1-specific modules associated with enriched GO terms highlighted in Fig. 3D**

UMAPs showing expression of the genes highlighted in Fig. 3D, demonstrating their enrichment in the cells assigned as "branch 1". Cells were grouped in hexagonal bins using schex[97].

Supplementary Figure 12

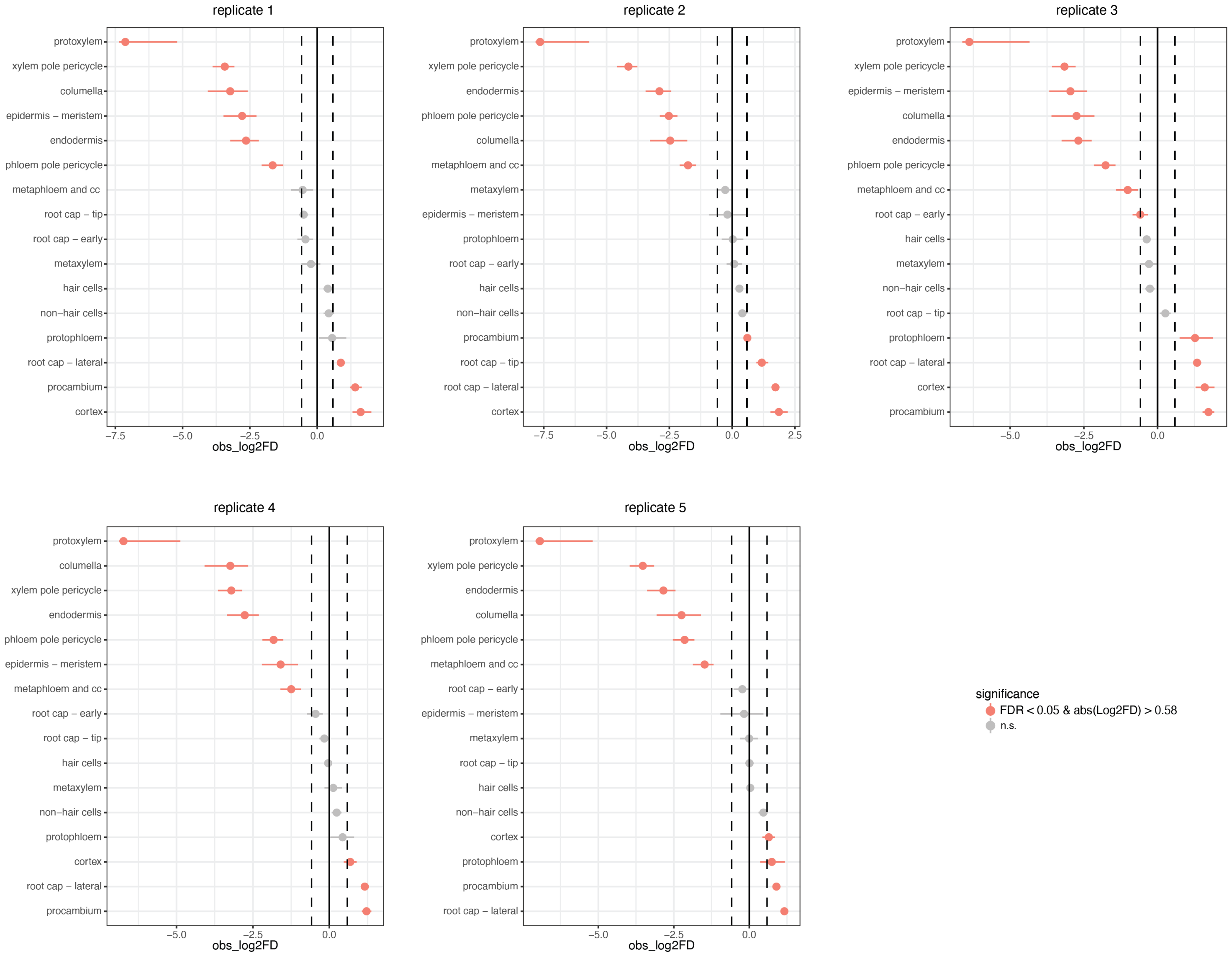

**Fig. S12. Changes in cell type abundance in *scr-3* root tips**

Point-range plot showing the confidence interval for the cell type proportional difference between the *scr-3* and Col-0 datasets, as compared to 1000 random permutations (see Methods). *scr-3* cells were compared to the five Col-0 replicates independently. Cell types whose proportion is significantly affected in *scr-3* are colored in pink (FDR<0.05 and |log2 fold difference|>0.58).

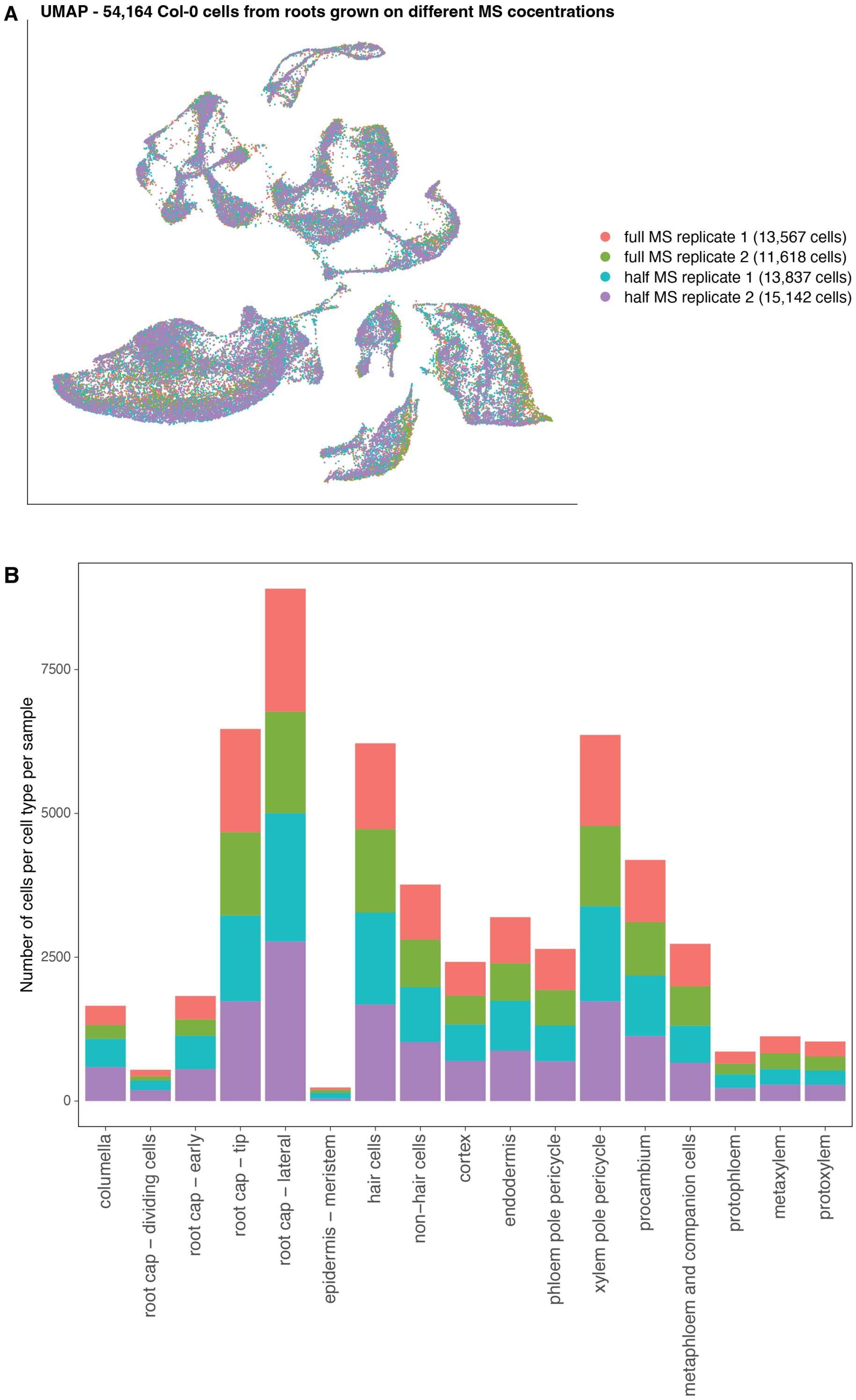
A
UMAP - 54,164 Col-0 cells from roots grown on different MS cocentrations
full MS replicate 1 (13,567 cells)
full MS replicate 2 (11,618 cells)
half MS replicate 1 (13,837 cells)
half MS replicate 2 (15,142 cells)
B
Number of cells per cell type per sample
0
2500
5000
7500
columella
root cap – dividing cells
root cap – early
root cap – tip
root cap – lateral
epidermis – meristem
hair cells
non–hair cells
cortex
endodermis
phloem pole pericycle
xylem pole pericycle
procambium
metaphloem and companion cells
protophloem
metaxylem
protoxylem

**Fig. S13. Samples under different nutrient concentrations have similar cell type compositions**

(**A**) UMAP showing the cells of the experiment testing the effect of nutrient concentration upon root tip gene expression states, described in Fig. 5E, colored by the sample they are derived from. (**B**) Barplot showing the number of cells per sample per cell type.

### Supplementary Figure 14

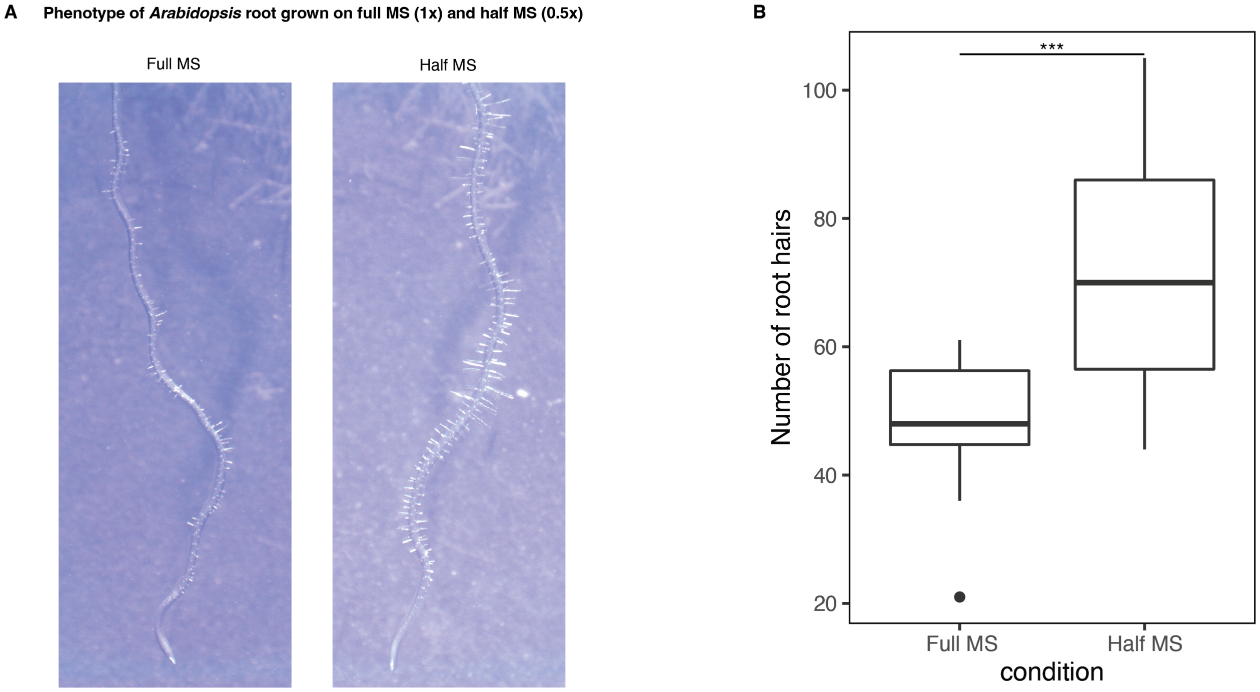

**Fig. S14. Phenotype of *Arabidopsis* roots grown on full MS (1x) and half MS (0.5x) media.**
(**A**) Images of roots grown vertically on full MS or half MS plates, 6 days after germination. (**B**) Boxplot showing the distribution of the number of root hairs per root in the first 2.5mm of roots grown on full MS or half MS. Asterisks indicate a p-value< 0.001 in the Wilcoxon Rank Sum test.

Supplementary Figure 15

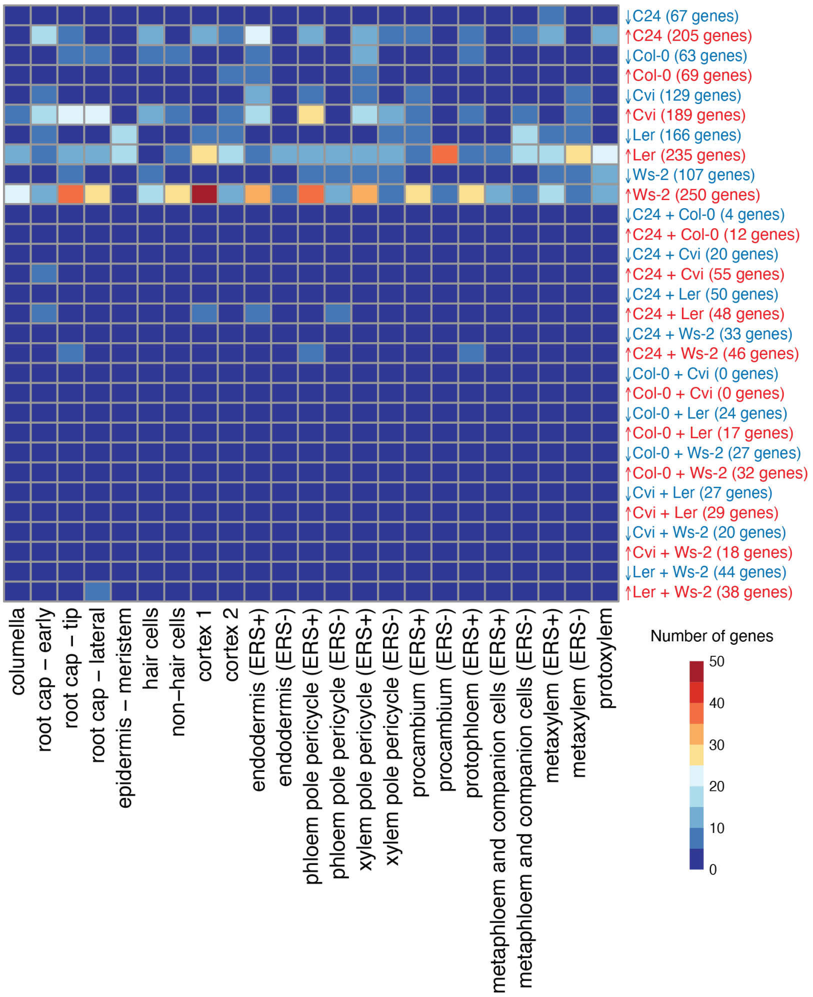

**Fig. S15. Accession-specific DEG are mainly specific to one accession rather than two**
Heatmap showing the number of genes found differentially expressed in one or two accessions as compared to others in the study, per cell state. The majority of genes exhibit differential expression specific to one of the accessions tested, rather than two.

**A Genes down-regulated specifically in C24**

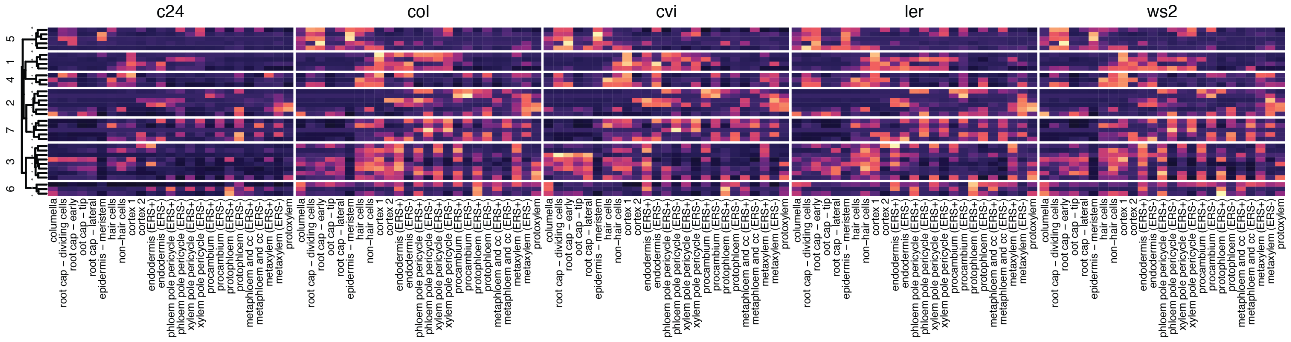


**B Genes up-regulated specifically in C24**

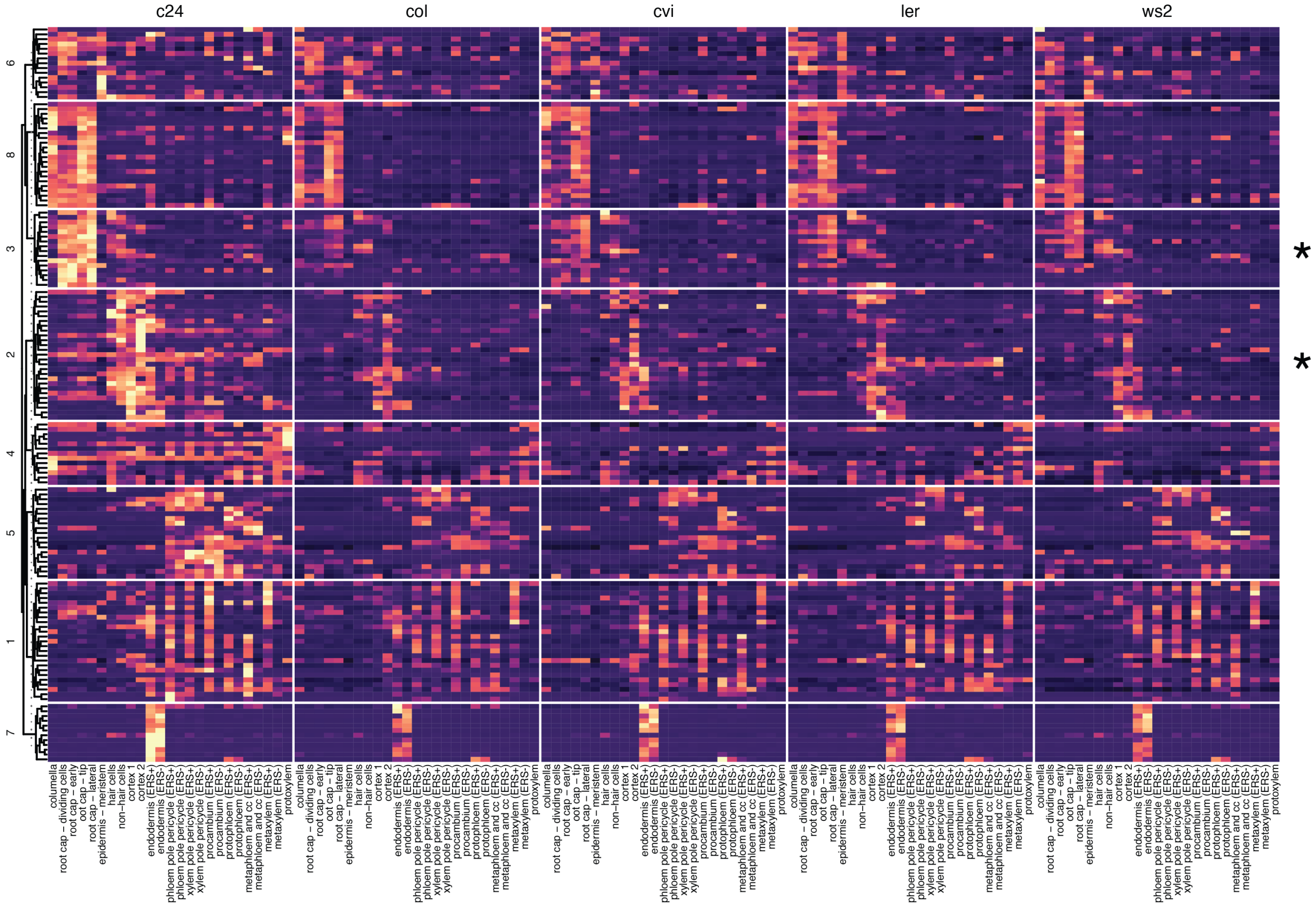


**C Genes down-regulated specifically in Col-0**

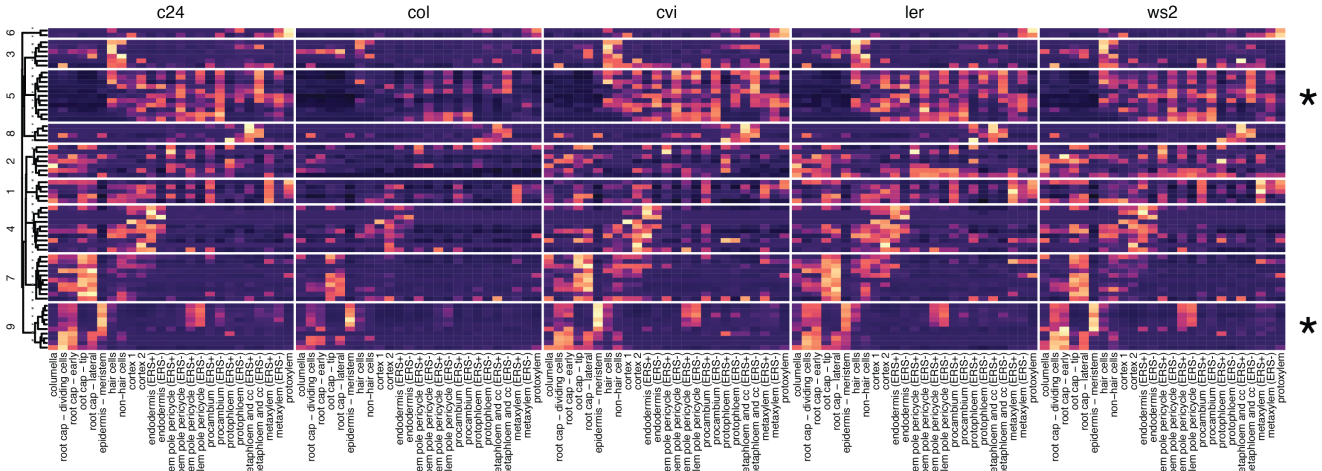

**D  Genes up-regulated specifically in Col-0**

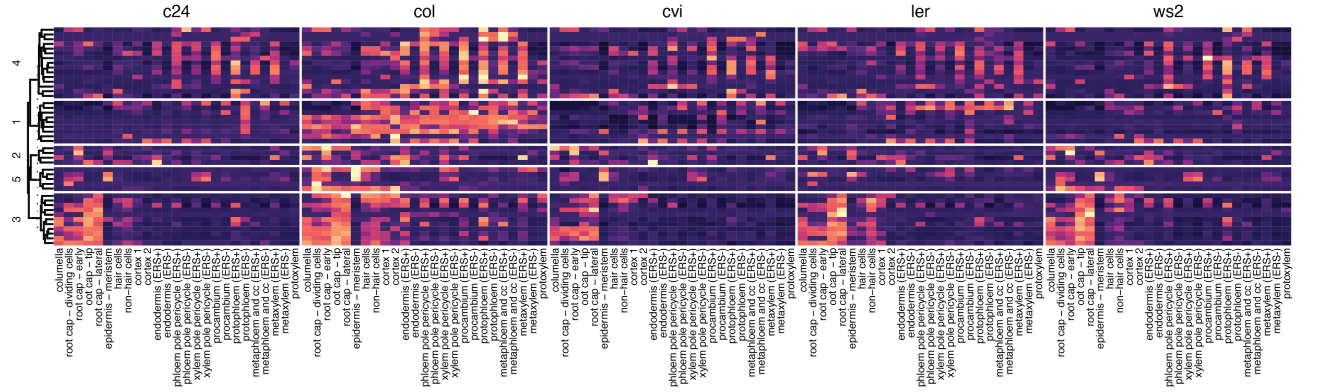


**E  Genes down-regulated specifically in Cvi**

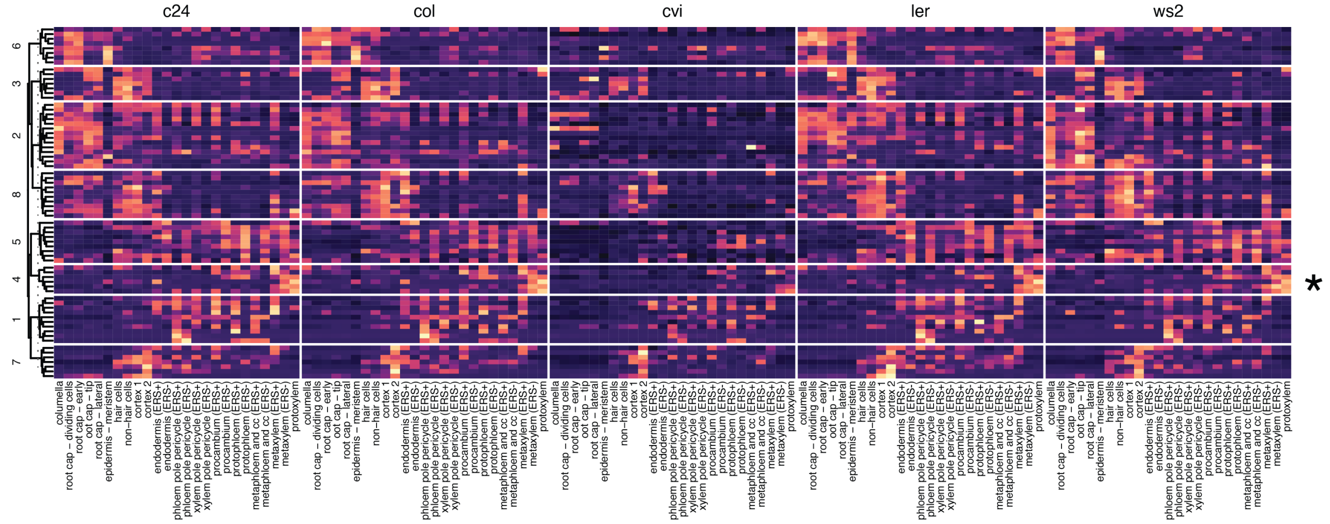


**F  Genes up-regulated specifically in Cvi**

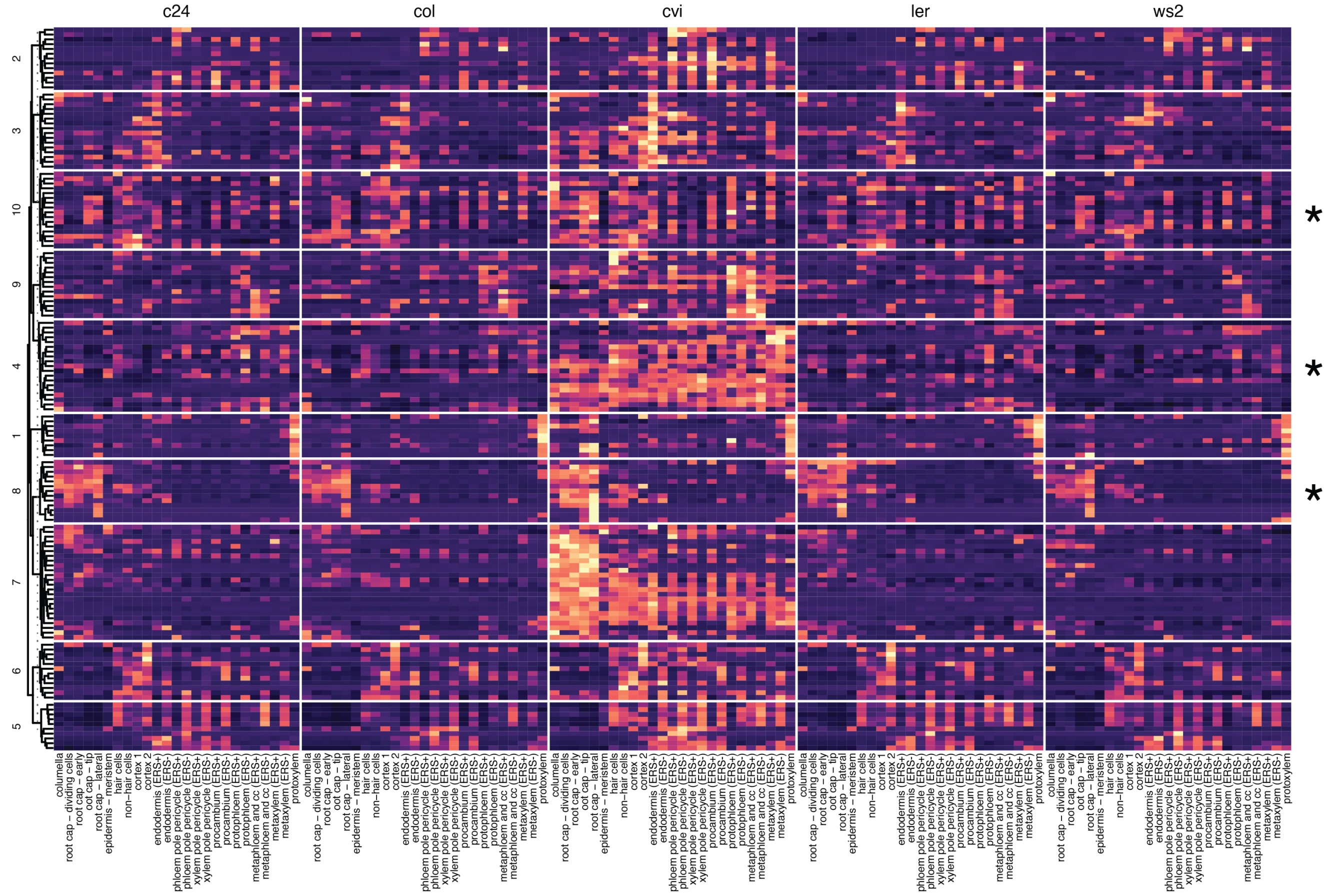

**G Genes down-regulated specifically in Ler**

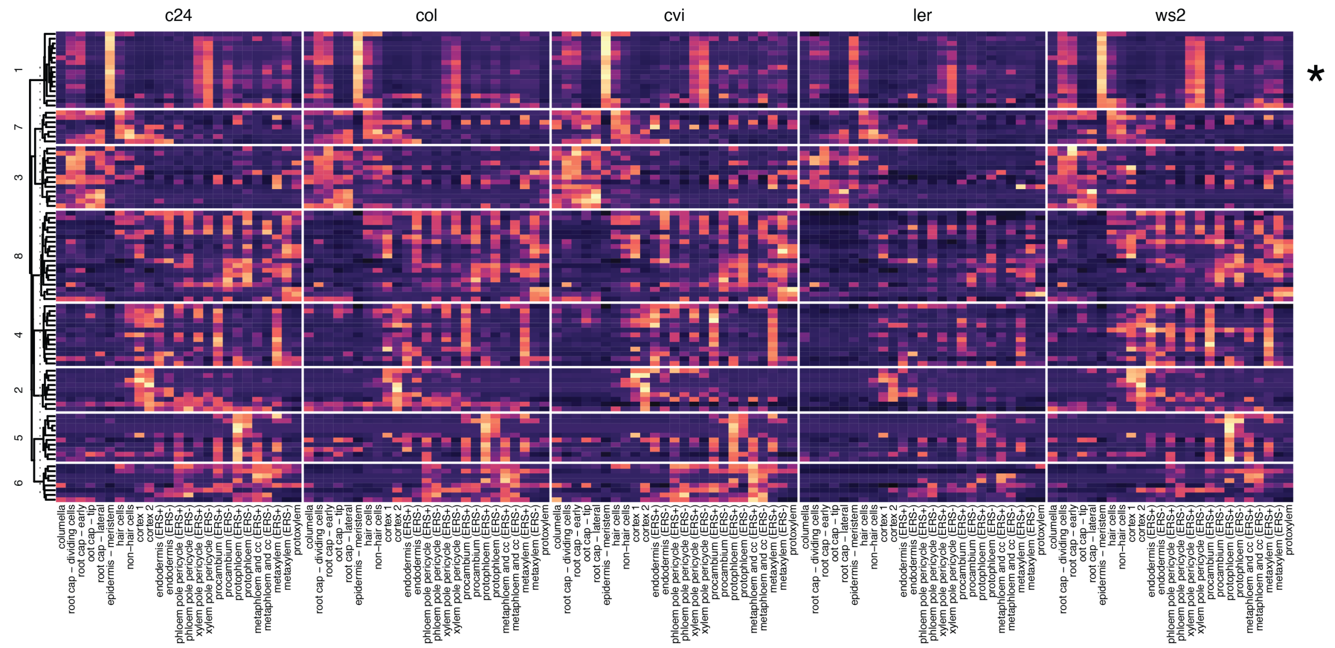


**H Genes up-regulated specifically in Ler**

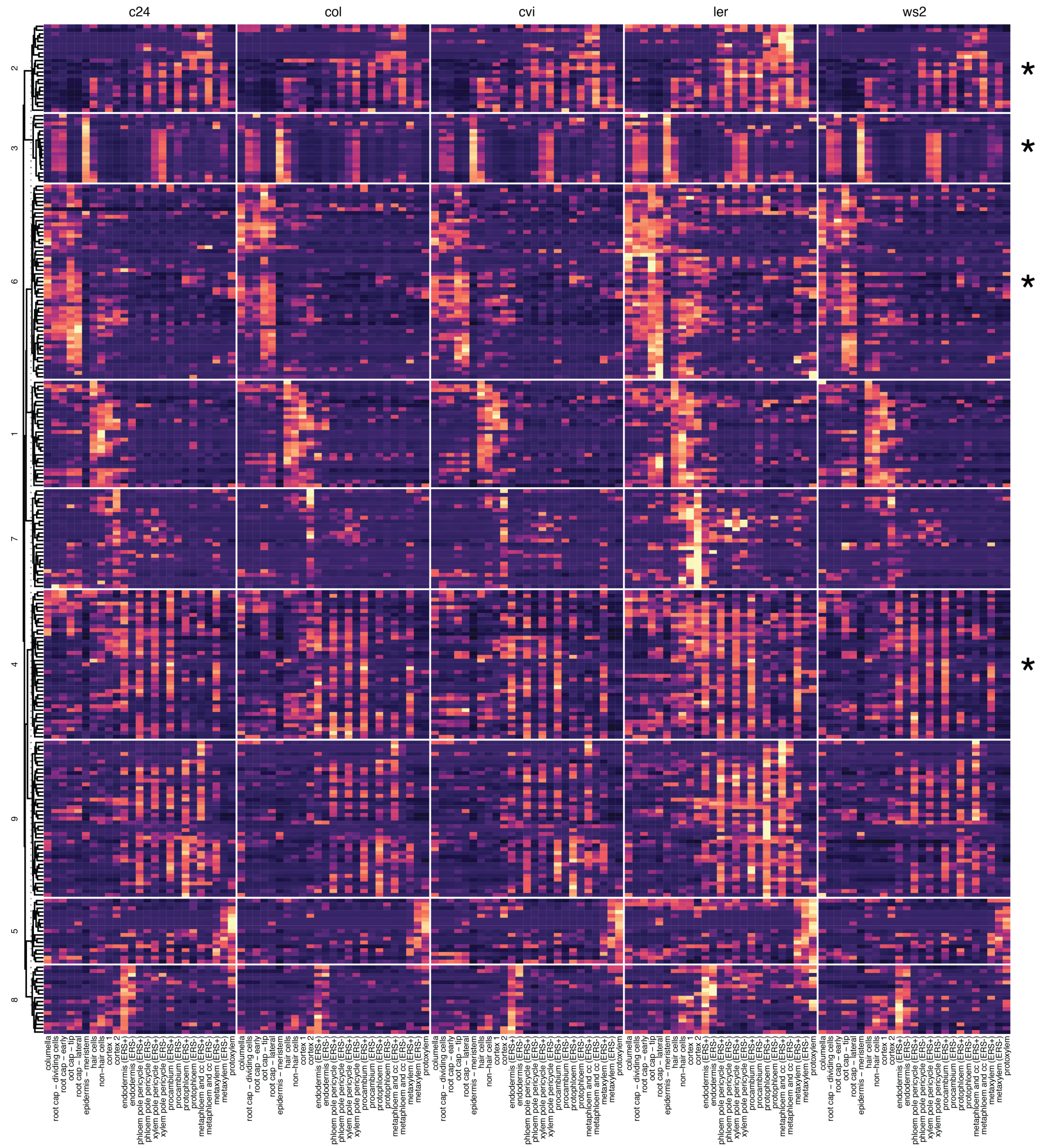

**I Genes down-regulated specifically in Ws-2**

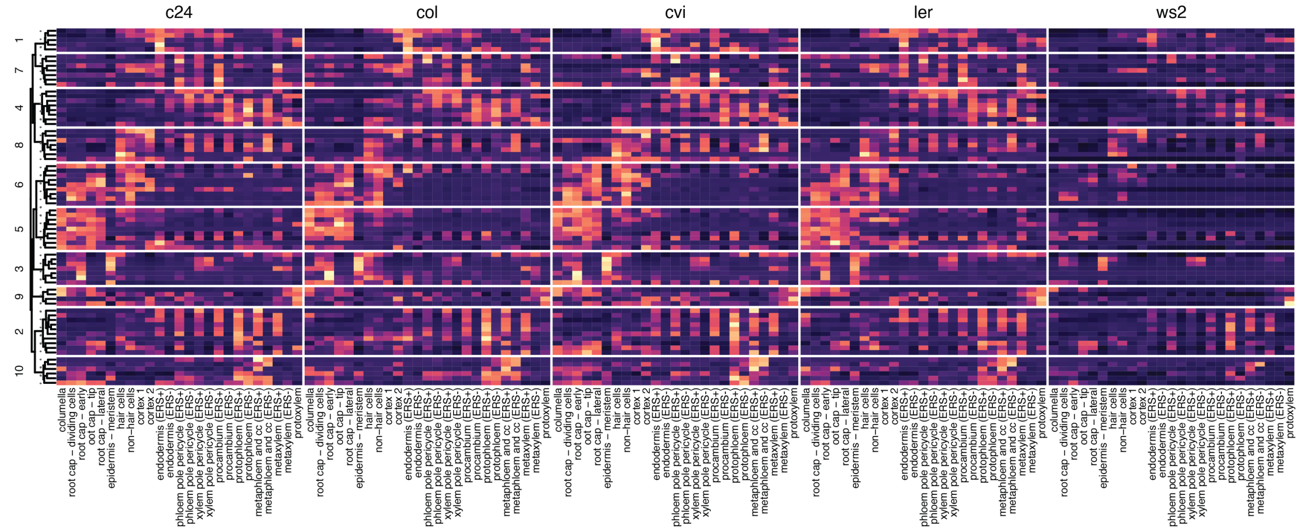


**J Genes up-regulated specifically in Ws-2**

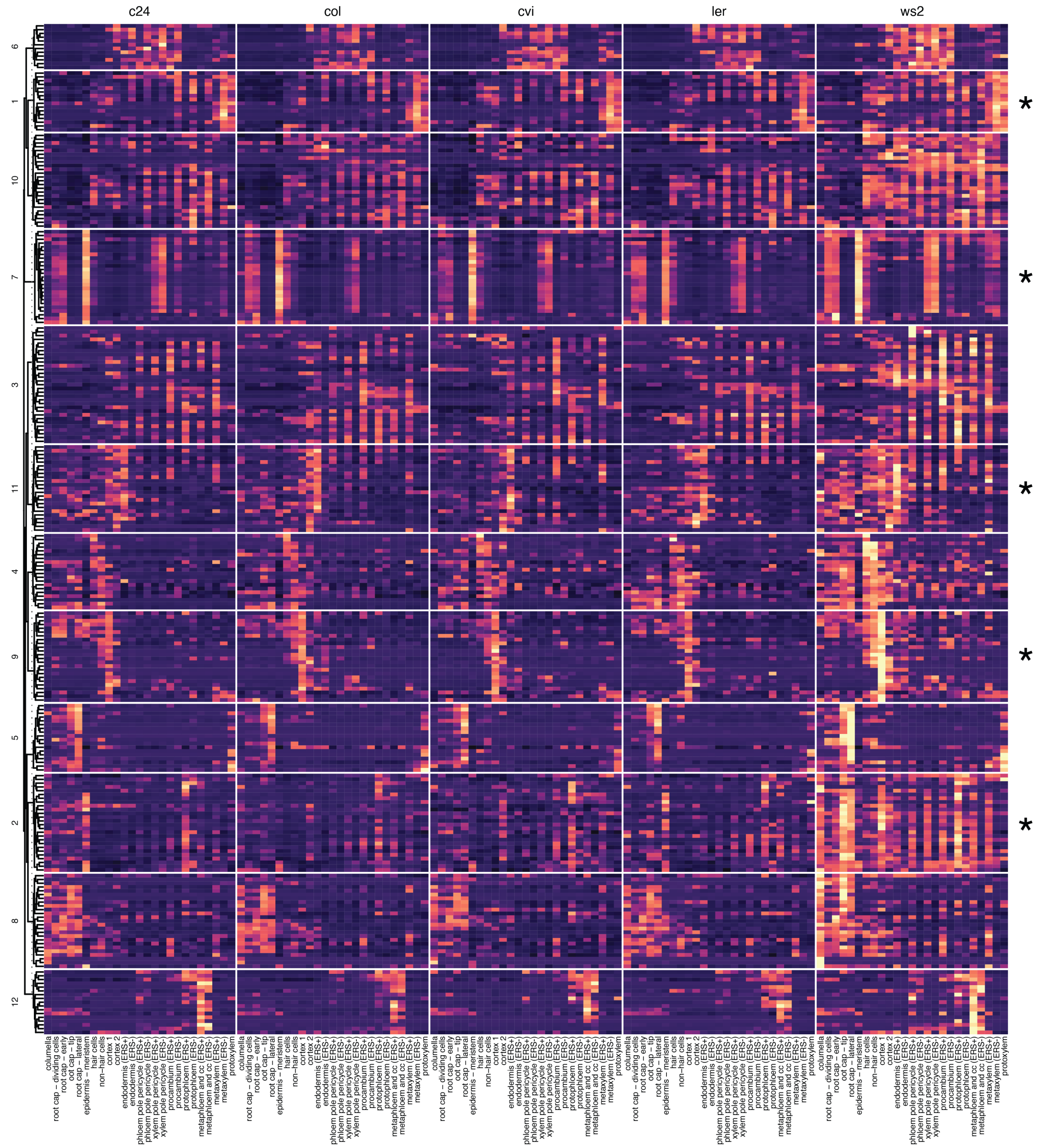

**Fig. S16. Accession-specific changes in cell states**

Heatmaps showing the scaled average expression of accession-specific DEGs, per cell state, per accession. In each category, genes were clustered based on their expression patterns. Asterisks indicate the eighteen clusters that have significant GO term enrichments, which are displayed in Fig. 6B.
