## Supplementary tables legend for "An environmentally responsive transcriptional state modulates cell identities during root development"

**Table S1. Genes affected by protoplast isolation**

AGIs of the genes that were identified as significantly up- or down-regulated during protoplast isolation (adjusted p-value <0.001 and a log2 fold change in expression >2)

**Table S2. Gene co-expression modules**

Composition of the 64 gene modules presented in Fig. 2A. Gene symbols and descriptions for each AGI was obtained from TAIR (www.arabidopsis.org)

**Table S3. Enrichment for putative ABA-regulated genes in co-expression modules**

Result of the enrichment test for genes that have a ABA-responsive element (ABRE) in the 500bp region upstream of their TSS according to the DAP-seq dataset (see Methods), per module. Modules that show a significant enrichment (padj < 0.05) are highlighted in orange.

**Table S4. Genes differentially expressed upon changes in nutrient availability**

Genes detected as significantly up- or down-regulated in plants grown on half MS as compared to full MS, and the cell type in which this differential expression was detected (t-test, padj <0.05).

**Table S5. Accession-specific differentially expressed genes**

Genes detected as significantly up- or down-regulated in one of the *Arabidopsis* accessions as compared to the other four tested, and the cell identity in which this differential expression was detected (t-test, padj <0.05).
